# Microbial methane cycling in a landfill on a decadal time scale

**DOI:** 10.1101/2023.01.20.524919

**Authors:** Daniel S. Grégoire, Nikhil A. George, Laura A. Hug

## Abstract

Landfills generate outsized environmental footprints due to microbial degradation of organic matter in municipal solid waste, which produces the potent greenhouse gas methane. With global solid waste production predicted to increase 69% by the year 2050, there is a pressing need to better understand the biogeochemical processes that control microbial methane cycling in landfills. In this study, we had the rare opportunity to characterize the microbial community responsible for methane cycling in landfill waste covering a 39-year timeframe. We coupled long term geochemical analyses to whole-community DNA (i.e., metagenomic) sequencing and identified key features that shape methane cycling communities over the course of a landfill’s lifecycle. Anaerobic methanogenic microbes are more abundant, diverse, and metabolically versatile in newer waste, fueling rapid methane production early in a landfill’s lifecycle. Aerobic methanotrophs were repeatedly found in leachate where low levels of oxygen were present and exhibited adaptations that aid survival under steep redox gradients in landfills. The potential for anaerobic methane oxidation, which has historically been overlooked despite anoxic habitats dominating landfills, was prevalent in a 26-year-old landfill cell which was in a state of slow methanogenesis. Finally, we identified the metabolic potential for methane oxidation in lineages that are widespread in aquatic and terrestrial habitats, whose capacity to metabolize methane remains poorly characterized. Ultimately, this work expands the diversity of methane cycling guilds in landfills and outlines how these communities can curb methane emissions from municipal solid waste.

**Significance:** Microbes are major contributors to methane emissions from solid waste however the temporal dynamics of methane cycling communities in landfills remain poorly understood. We addressed this gap by using whole-community DNA (i.e., metagenomic) approaches to characterize microbial methane cycling in waste covering a 39-year timeframe. We show that methane-producing microbes are more abundant, diverse, and metabolically versatile in new waste compared to old waste. We highlight that methane oxidation in the absence of oxygen is overlooked in landfill biogeochemical models and that novel lineages can potentially contribute to methane sinks across a broad range of habitats. These findings can strengthen predictive models for methane cycling in landfills and inform sustainable waste management strategies to curb methane emissions from solid waste.

## Introduction

Landfills are one of the key mitigation gaps in managing global methane emissions. From 2000 to 2017, landfills produced 60 to 69 Tg of methane per year (1, 2). In high-income countries such as the United States, landfills account for up to 20% of net methane emissions (3). As of 2018, it is estimated that 2.01 billion tonnes (i.e., 2010 Tg) of solid waste has been produced globally, with 35% of this waste being landfilled (4). With solid waste production predicted to increase to 3.40 billion tonnes (i.e., 3400 Tg) by 2050 (4), there is a pressing need to better understand the biogeochemical processes that control methane’s fate in landfills to enable development of waste management practices that mitigate greenhouse gas (GHG) emissions.

Landfilled waste is spatially and geochemically heterogeneous, with compositional changes occurring over extended time scales. Part of the challenge in managing GHG emissions from municipal solid waste (MSW) lies in predicting how these variables interact to affect methane cycling over decades of landfill operation.

The major biogeochemical transitions that occur in a sanitary landfill can be summarized using a five phase conceptual model [(5, 6) and references therein]. Phase 1 is the **aerobic phase**, where chemoheterotrophic microbes consume oxygen to metabolize organic carbon from paper, food waste, and cover soils. Phase 2 is the **anaerobic acid phase**, where fermentative microbes hydrolyze cellulose-bearing waste and produce labile organic substrates that support fermentation and organic acid production, which decreases pH in the landfill. Phase 3 is characterized by **rapid methanogenesis**, where labile organic and inorganic carbon substrates stimulate biogenic methane production by anaerobic archaea. Phase 4 is delineated by a transition to **slow methanogenesis**, where substrates that support methanogenesis have been depleted and methane production slows. In phase 5, **oxygen infiltration** can occur because substrates that support aerobic heterotrophy have been exhausted, such that aerobic respiration cannot outpace oxygen diffusion from the atmosphere. Phase 5 is considered to be the point at which MSW has stabilized yet remains the least well understood of the lifecycle phases because it can take over 20 years to develop and there are limited datasets covering this timespan.

Microbial metabolisms drive every major biogeochemical change that occurs in a landfill. A key 16S rRNA amplicon sequencing survey that sampled 19 landfills in the United States suggested that the age of refuse and local environmental conditions play significant roles in shaping microbial communities in landfill leachate (7). Several smaller-scale amplicon sequencing studies have suggested that variables including age (8), nutrient concentrations (9), physicochemical parameters (e.g., temperature and pH) (10), and contaminant concentrations (11–13) all shape microbial community structure in landfills. 16S rRNA surveys have also been used to characterize the succession of microbial taxa over the course of waste degradation in controlled settings (14–17). Methane cycling guilds in landfills have been examined using 16S rRNA primers specific to methanogens and methanotrophs (18, 19). Sequencing the *mcrA* gene, which codes for the methyl coenzyme M reductase responsible for converting methyl coenzyme-M to methane during methanogenesis (20), has expanded the diversity of methanogenic taxa associated with landfills (21, 22). Similarly, sequencing the *pmoA* gene, which codes for a key subunit of the particulate methane monooxygenase (pMMO), and the *mmoX* gene, which codes for a key subunit in the soluble methane monooxygenase (sMMO), has clarified the structure of methanotrophic communities in landfill cover soils (23–25). These studies highlight how the structure of microbial communities in a landfill can change over the course of biogeochemical transitions that control methane cycling.

Although amplicon sequencing surveys have shed light on the biodiversity contained within landfills, they have provided a limited understanding of the major guilds and physiological pathways controlling biogeochemical cycles in landfills. In the case of methanogens, taxonomy is routinely used to infer whether hydrogenotrophic (i.e., H_2_ and CO_2_ requiring) or acetoclastic (i.e., acetate requiring) methanogenesis contributes to methane production. This approach is limited when applied to novel methanogenic taxa that are not related to well-characterized model organisms and tends to ignore the contributions of less-well studied methanogenesis pathways (e.g., methylotrophic methanogenesis). Taxonomy is also used to determine whether methane oxidation is carried out by bacteria that require low levels of methane and high levels of key trace nutrients (i.e., type I methanotrophs) or high levels of methane but low levels of other key substrates (i.e., type II methanotrophs) (26). The use of these classifications has been called into question based on large-scale surveys of the *pmoA* gene, which revealed that methanotrophs from a broad range of habitats exhibit more diverse lifestyles than can be captured by the dichotomy indicated above (27). The potential for methanotrophy via mechanisms that do not require exogenous oxygen (e.g., intra-aerobic or anaerobic methane oxidation) are also rarely considered despite landfills being dominated by anoxic habitats that could support such pathways.

The recent application of metagenomic sequencing to landfills offers a promising solution to the limitations of previous work. Metagenomics has allowed valuable insights into the physiological pathways contributing to waste degradation including cellulose metabolism (28, 29) and plastic biodegradation (30). Metagenomics has also been used to address human health concerns tied to landfills such as the occurrence of antibiotic resistance in pathogens found in MSW (31, 32). Genome-resolved metagenomics (33) has the potential to connect methanogenic lineages to the substrates that can support methane production within a given landfill environment (34). Similar approaches can be used to develop metabolic models for methanotrophs and shed light on the range of methanotrophic lifestyles present in landfills (e.g., facultative vs. obligate, aerobic vs. microaerophilic vs. anaerobic). To date, metagenomic surveys that span the spatial and temporal scales relevant to landfill lifecycles are lacking.

We used metagenomic sequencing to provide an unprecedented historical perspective on microbial methane cycling in a sanitary landfill. In this study, we compare methanogen and methanotroph community structure and metabolic capacity in leachate samples from a landfill spanning the five landfill lifecycle phases. We use phylogenomic analyses and metabolic models to identify adaptations in methanotrophs and expand the diversity of taxa potentially capable of oxidizing methane in oxygen-limited landfill habitats. Finally, we demonstrate how the biodiversity in landfills includes and allows identification of microbes with uncharacterized methane cycling capabilities whose role in the global methane cycle has been overlooked.

## RESULTS AND DISCUSSION

### Landfill lifecycle geochemistry

The site sampled in this study is a sanitary landfill located in the northeastern United States (anonymity by request of site management). The site is equipped with leachate collection and biogas capture systems. MSW at the site is housed in six landfill cells, referred to herein as A, B, C, D, E, and F, which had operated in succession for 39 years at the time of leachate sampling in February 2019. A is the oldest cell (receiving waste from 1980 to 1982) and F is the youngest, receiving waste as of 2014. Landfill cells A, B, and C are closed and completely capped from receiving waste. Cells D, E, and F remain partially capped due to the implementation of a phased landfilling approach that will eventually create one contiguous landfill cell. The practice of leachate recirculation has shifted over time as MSW management strategies at the landfill have changed, such that leachate from any landfill cell could be recirculated through the parts of the landfill actively receiving waste. At the time of our sampling, leachate recirculation across the entire landfill site had stopped, which would limit the capacity of older landfill cells to affect the leachate geochemistry of newer ones receiving waste. To facilitate comparisons within such a heterogeneous system, we organized our geochemical analyses around the most active filling periods for each landfill cell.

A total of 8 samples were obtained as part of this campaign [referred to as A, B, C, D1, D2, E, F1, and F2]. Samples D1 and D2 denote leachate collected from two different wells associated with different parts of landfill cell D. These two samples are associated with different drainage areas in cell D that began receiving waste in 1993 and 1995, respectively. The drainage area associated with sample D1 receives leachate from cell D, but also from a valley between cells C and D where the leachate from both cells mixes and where gas is monitored via dedicated sampling ports. Samples F1 and F2 denote leachate collected from two different locations in cell F, which began receiving waste in 2014 and 2016, respectively. Cell F was actively receiving waste at the time of sampling, as were parts of cell D where the cap was removed from areas of cell D that abutted with cell F. Cell E was largely capped at the time of sampling except for the southeast face which is comprised of soil and vegetative cover. Despite these connections, distinct trends in leachate geochemistry were observed at the local scale for each landfill cell. We have used the age of MSW to ground our classification of each cell into the biogeochemical phases of a landfill lifecycle.

#### Landfill cells A, B, and C (A: filling from 1980-1982, 39 years old, capped; B: filling from 1982-1988, 37 years old, capped; C: filling from 1988-1993, 31 years old, capped)

Landfill **cells A and B can be classified to phase 5** of the landfill lifecycle whereas **cell C is in phase 4**. These landfill cells passed their peak chemical oxygen demand (COD) and biological oxygen demand (BOD) after ~5 years of operation, maintain low redox potentials (i.e., between −200 to 200 mv), and have low concentrations of carbon substrates that could support methanogenesis (i.e., <10 mg L^-1^ for organic acids and decreasing levels of bicarbonate) (**Fig 1** and **Fig S1**). The median values for total gas flared from cells A, B, and C are considerably lower compared to newer landfill cells such as E and F (i.e., 20 to 100 vs. 400 to 530 standard cubic feet per minute respectively), suggesting methane production has slowed in the oldest parts of the landfill (**Fig 2A**). Oxygen consistently accounts for ~5 % of the gas flared from A and B but is <1 % of the gas flared from cell C (**Fig 2B**). Although data from a gas flaring vent located in the valley that received waste between the filling period for cells C and D (i.e., 1998 to 1999) suggests some oxygen may be infiltrating cell C (see **Fig 2B**), we opted to classify cell C to phase 4 because it lacked the stronger evidence of oxygen intrusion seen for cells A and B.

**Figure 1:**
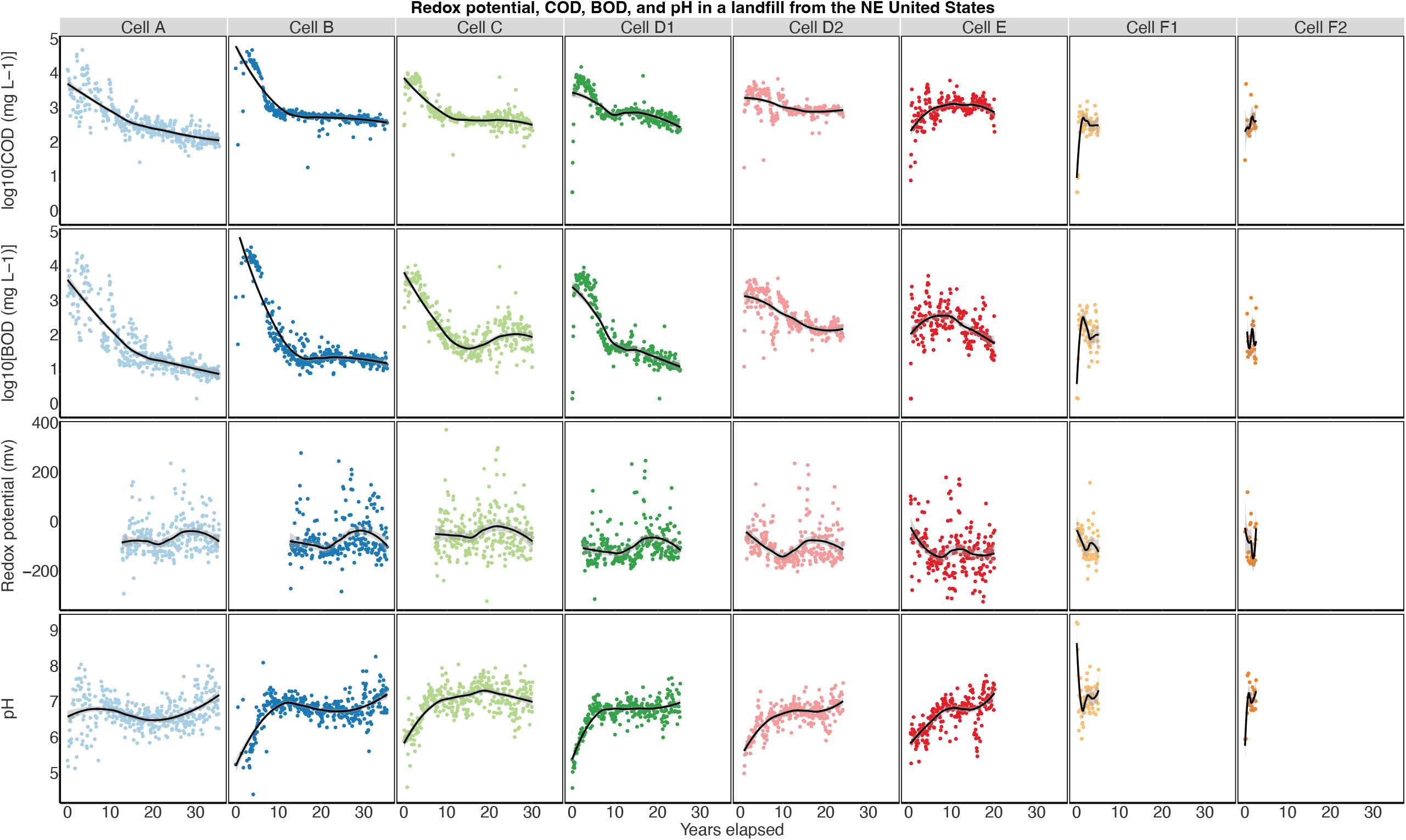
Aqueous geochemistry of landfill leachate collected from landfill cells A, B, C, D1, D2, E, F1 and F2 from 1983 to 2019. These data were compiled from monitoring records provided by the site management. Loess curves are presented to visualize smoothed temporal trends. Abbreviations denote the following: COD = Chemical oxygen demand, and BOD = biological oxygen demand.

**Figure 2:**
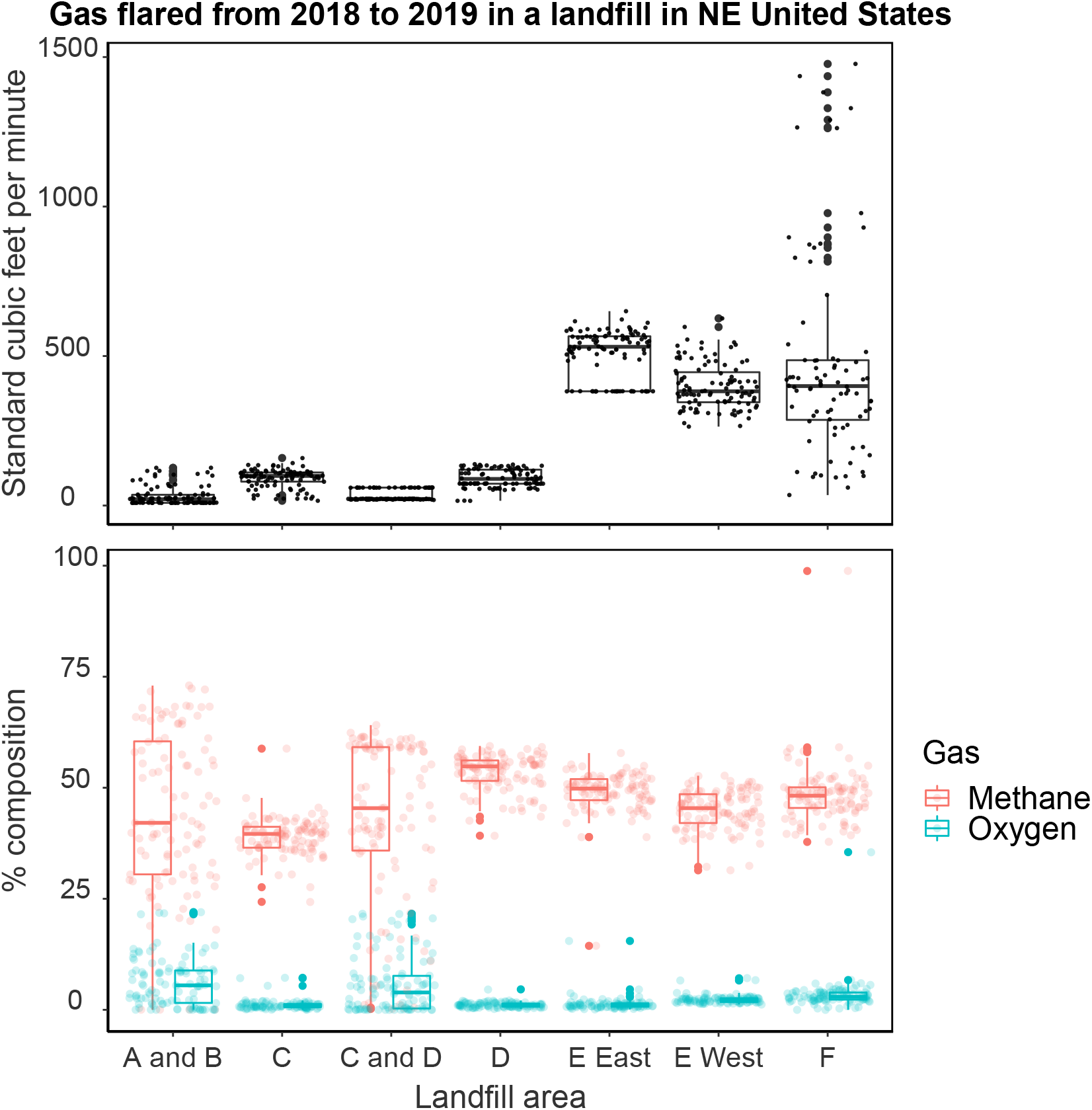
Total gas flared (upper panel) and the relative composition of the gas flared (bottom panel) from vents associated with landfill cells A, B, C, D, E, and F. This data was provided by the site owners for sampling years 2018-2019 to cover seasonal variation over two years and capture current trends in gas emissions rather than historical ones.

#### Landfill cell D (D1: filling from 1993-1998, 26 years old, D2: filling from 1995-1998, 24 years old)

Both sampling locations in cell D suggest this part of the landfill has entered **phase 4 of slow methanogenesis**. Both locations at cell D passed their peak COD and BOD after ~5 years of operation as seen for cells A, B, and C (see **Fig 1**). Historical peaks in organic acids are higher for D1 vs D2, however both locations experienced peaks and subsequent decreases in organic acids followed by decreases in bicarbonate and increases in pH over the same timeframes as cells A, B, and C (**Fig 1** and **Fig S1**). The median value for total gas flared from location D1, which is connected to the C/D drainage area separate from location D2, is lower compared to location D2 connected to drainage area D (i.e., 22 vs 89 standard cubic feet per minute, see **Fig 2A**). The gas flared from D2 is largely comprised of methane with oxygen consistently accounting for <1% of the gas composition in this cell (**Fig 2B**). In the case of location D1, there is some evidence of oxygen infiltration based on the flaring data from the C/D valley vent (see **Fig 2B**). We applied the same logic as we did for cell C, where the weak evidence of oxygen intrusion considered alongside the younger age of D1 vs. cells A and B resulted in the entirety of cell D being classified to phase 4 of slow methanogenesis.

#### Landfill cell E (filling from 1999 to 2014, 20 years old)

Based on the geochemistry of the leachate and gas flaring data, **cell E was classified as being in phase 3 of rapid methanogenesis**. Landfill cell E reached peak COD and BOD over a similar timeframe compared to cell D and pH has increased steadily from ~6 to 7.8 while maintaining reducing conditions (i.e., redox potential consistently around −200 mV), suggesting cell E has moved past the anaerobic acid phase (**Fig 1**). The peaks in organic acid concentrations in cell E are less pronounced compared to the landfill cells A-D, suggesting historically lower inputs of substrates that support fermentation and the production of organic acids (**Fig S1**). Indeed, a waste diversion program diverting yard waste was implemented in the year 2000, which likely resulted in lower volumes of organic matter deposited into cell E (communication from site owners).

These results could also be attributed to the connection between landfill cell D and E, wherein portions of cell D remained uncapped during landfilling of cell E to redistribute moisture from leachate across both landfill cells (communication from site owner). Lower moisture in cell E would have limited the circulation of nutrients and the maintenance of anoxic habitats required for fermentative acid production such that organic acid concentrations decreased. Cell E also experienced a decline in bicarbonate concentrations, reaching concentrations as low as 851 mg L^-1^ before increasing to concentrations comparable to the peaks observed for landfill cells A, B, C, and D (i.e., ~3500 mg L^-1^) (**Fig S1**). We make note of this bicarbonate depletion because it was observed on 2019-02-12, the day of our sampling trip, which may have impacted the observed microbial community.

Gas flaring data for cell E show total gas production is almost an order of magnitude higher compared to cells A, B, C, and D, (i.e., 531 and 399.50 standard cubic feet per minute for E East and E West, respectively) demonstrating higher methane production for cell E (**Fig 2A**). The composition of the gas coming from cell E is nearly identical at both locations, with oxygen accounting for 1-2% of the gas flared (**Fig 2B**). These observations suggest that environmental conditions are homogenous enough in cell E to support relatively uniform methane production. Methane production in cell E does not seem to be impacted by notable decreases in inorganic carbon in the leachate or physical connections to cells D and F.

#### Landfill cell F (F1: filling from 2014-present, 5 years old, F2: filling from 2016 to present, 3 years old)

Cell F is classified as **transitioning from phase 2 of anaerobic acid production to phase 3 of rapid methanogenesis** based on contrasting leachate geochemistry from two sampling locations. Unlike other locations in the landfill, location F1 recently experienced an increase in COD alongside a decrease in BOD, whereas location F2 showed a steep increase in both COD and BOD (**Fig 1**). These observations suggest an influx of organic carbon that can be oxidized, in line with phase 2 of anaerobic acid production, although the pH remains circumneutral (see **Fig 1**). These increases in COD and BOD are accompanied by small peaks in organic acid production alongside rising bicarbonate concentrations, which may have offered strong buffering that limited decreases in pH in this part of the landfill (**Fig S1**). These increases in bicarbonate are in line with a habitat that can still support heterotrophy and can be attributed to the mineralization of MSW.

The total amount of gas produced at the monitoring location for cell F is comparable to both locations monitored at cell E (**Fig 2A**). In contrast to cell E, which we classified as being in a state of rapid methanogenesis, there was more variability in the amounts of gas produced at cell F with maxima reaching > 1000 standard cubic feet per minute and minima as low as 35, with median oxygen composition being low at ~ 2% (**Fig 2**). The depletion in organic carbon alongside variable amounts of gas being flared with a consistent methane composition is in line with what we would expect of a habitat where the microbial community is beginning to initiate methane production, where methanogens are competing with heterotrophs for labile substrates.

### Methanogen community

From the initial 1,881 metagenome-assembled genomes (MAGs) obtained from landfill metagenomes (see **Methods**, and **Supplemental Table 1** for metagenome statistics), 74 MAGs were identified as putative methanogens coming from leachate samples. The putative methanogen MAGs were taxonomically classified into 3 phyla following curation: *Halobacterota, Thermoplasmatota*, and *Euryarchaeota* and 11 families (**Fig 3**). Two MAGs could not be classified to the family level: STE_86, classified to the order *Methanobacteriales*, and STF1_149, the only MAG classified to the order *Methanofastidiosales* (see **Fig 3** and **File S1**).

**Figure 3:**
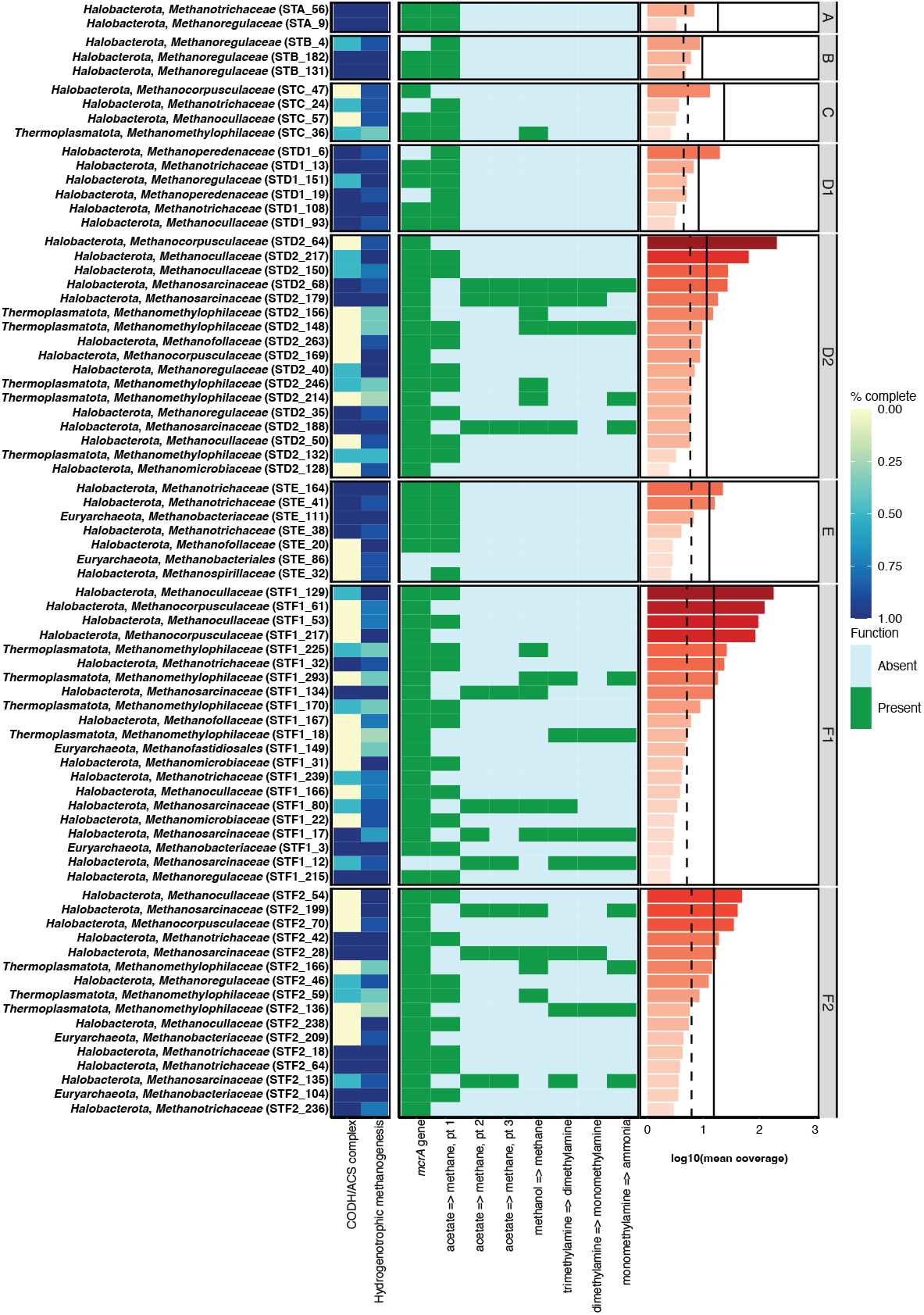
Metabolic heatmap and coverage data for putative methanogenic MAGs. The abbreviation CODH/ACS complex denotes the completion of the carbonic anhydrase/acetyl-CoA synthase pathway. Coverage values denote the mean coverage of all scaffolds for a putative methanogenic MAG under a log_10_ transformation. Solid black vertical lines denote the mean coverage calculated for MAGs at the whole-community level and dashed black lines denote the median coverage calculated for MAGs at the whole-community level.

#### Landfill cells A, B, and C

The putative methanogen community from these areas of the landfill displayed low diversity and low abundance, in line with our characterization of these habitats as being in phase 5 (cells A and B) and phase 4 (cell C) based on geochemistry. Landfill cells A, B, and C contained two, three, and seven putative methanogen MAGs, respectively. In all cases, putative methanogen MAGs had coverage values close to the median coverages for their respective datasets and were an order of magnitude or more below the coverages of the most abundant MAGs in each landfill cell metagenome (see **Fig 3, Fig S2**, and **File S1**).

All putative methanogen MAGs obtained from landfill cells A and B can be classified as acetoclastic methanogens based on the presence of the acetyl-CoA synthetase (i.e., noted as acetate => methane pt.1 by DRAM), high completion (i.e., > 50 %) of the carbon monoxide dehydrogenase/acetyl-CoA synthase (CODH/ACS) complex supporting acetyl-CoA dismutation, and high completion (i.e., > 75%) of the hydrogenotrophic methanogenesis pathway (**Fig 3**). The functional potential for methanogenesis from cell C shows a mix of strictly hydrogenotrophic, acetoclastic, and methylotrophic methanogens (see **Supplemental Results** and **Methods**).

When considering the microbiological analyses alongside our predictions based on geochemistry, the low abundance and diversity of methanogens are in line with the observed low rates of methane production in these ageing landfill cells. The restricted metabolic pathways that can support methanogenesis align with observations of these habitats having limited substrates available that can support methanogenesis. Furthermore, the lower diversity and abundance of methanogens in cells A and B could also be attributable to oxygen infiltration, which would inhibit methanogenic metabolism (**Fig 2**).

#### Landfill cell D

### Landfill cell D appears to be in phase 4 of slow methanogenesis

Looking at the microbial community from locations D1 and D2, we see contrasting trends in the putative methanogenic community structures that suggest this landfill cell is not experiencing a uniform transition from phase 3 to phase 4. The community from D1 is comprised of six MAGs, and more closely resembles that of cells A, B, and C than D2. Many of the families present in D1 were detected in cells A, B, and C, including the *Methanotrichaceae, Methanoregulaceae*, and *Methanocullaceae* (**Fig 4**). Two MAGs from the *Methanoperedenaceae* were also recovered from D1 based on detection of *mcrA* genes (i.e., STD1_6 and STD1_19) (**Fig 3**). The detection of the *Methanoperedenaceae* is noteworthy as they are the only named family of anaerobic methane oxidizing Archaea (ANME), as such, these MAGs were not included in the initial total count of putative methanogens. The MAGs belonging to the *Methanoperedenaceae* are discussed within the examination of methanotrophy, below.

**Figure 4:**
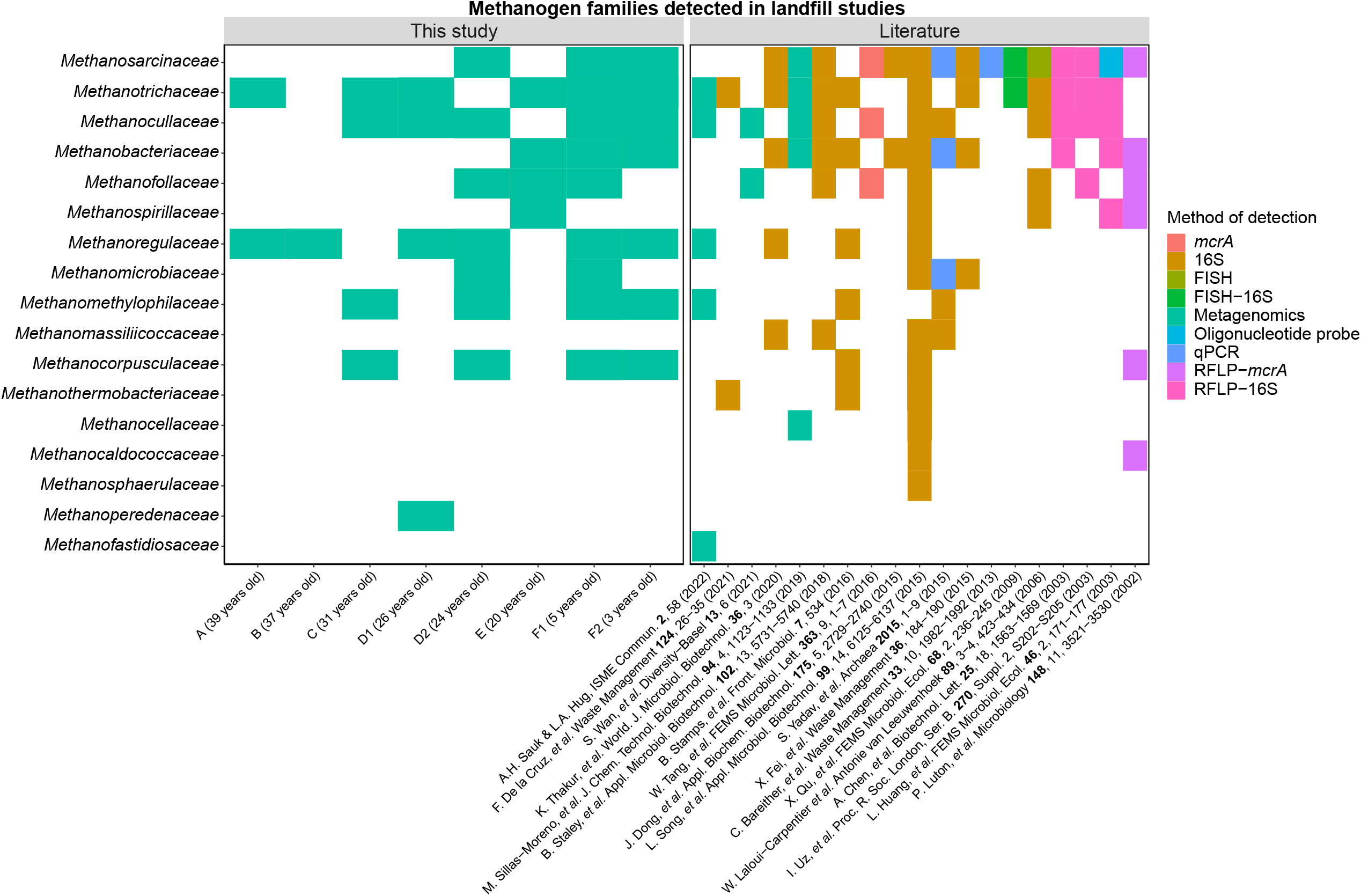
Methanogen and *mcr*-bearing families detected in this study and previously published work using different analytical techniques. Where possible, all taxonomic labels have been updated to reflect the GTDB r89 taxonomy used in our study. Abbreviations denote the following: 16S (shorthand for 16S rRNA amplicon sequencing), FISH (fluorescent *in situ* hybridization), RFLP (restriction fragment length polymorphism), and qPCR (quantitative polymerase chain reaction). Abbreviated references have been provided and the raw input data used to generate this figure can be found in **File S4**. Data obtained from (in chronological order): (7, 11, 14, 15, 17–19, 21, 22, 32, 34, 49, 88–93).

In contrast to location D1, 17 putative methanogen MAGs were recovered from location D2. There was considerable overlap between the families detected in D2 and cells A, B, C, and location D1 (**Fig 4**), with three additional families identified at location D2 (**Fig 4**).

In direct contrast to trends of low abundance for putative methanogens in older landfill cells, MAG STD2_64 from the family *Methanocorpusculaceae* displayed the maximum coverage value for all MAGs assembled from location D2 (i.e., 205.95; see **Fig 3, Fig S2**, and **File S1**). Additional MAGs from the families *Methanocullaceae* (e.g., STD2_217 and STD2_150) and *Methanosarcinaceae* (e.g., STD2_68 and STD2_179) had coverage values ranging from 18.25 to 64.71, which were higher than the median and mean coverage for the whole community (i.e., 5.85 and 11.50, respectively; **Fig 3**).

The differing trends observed at cells D1 and D2 extend to the methanogenesis pathways predicted at both locations. The capacity for methanogenesis at D1 resembles that of cells A and B, with MAGs from the *Methanotrichaceae, Methanoregulaceae*, and *Methanocullaceae* all classified as acetoclastic methanogens based on the presence of genes coding for the acetyl-CoA synthetase, >50 % complete CODH/ACS complexes, and >75 % complete hydrogenotrophic methanogenesis pathways (**Fig 3**). More substrates can potentially support methane production at D2. Abundant MAGs from the *Methanocorpusculaceae, Methanofollaceae, and Methanomicrobiaceae* were classified as strictly hydrogenotrophic methanogens (see **Methods** and **Supplemental Results** for details). Acetoclastic methanogens were identified from two families (*Methanocullaceae* and *Methanoregulaceae*), and strict methylotrophic methanogens were classified to the family *Methanomethylophilaceae*. The broadest capacity for methanogenesis was observed in MAGs from the *Methanosarcinaceae* family (i.e., STD2_68, STD2_179, and STD2_188), which displayed the potential for hydrogenotrophic, acetoclastic, and methylotrophic methanogenesis (**Fig 3**).

Despite initially classifying both locations in landfill cell D as being in phase 4 of slow methanogenesis, our microbiological observations suggest biogeochemical succession is occurring in a more segregated fashion. Although location D1 houses waste that is only 2 years older than location D2, the community structure and methanogenic capacity observed at D1 is more in line with landfill cells that are 10 years older. This discrepancy may be occurring due to cell D’s connection to the older cell C. In D2, methanogens are the most abundant populations and methanogenesis is predicted to use a broader array of substrates, suggesting substrates capable of supporting methanogenesis may be more widely available at D2 compared to D1.

Concentrations of inorganic carbon were higher at D2 compared to D1 in the timeframe surrounding our sampling expedition (i.e., ~2500 to 3500 mg L^-1^ vs ~ 1000 mg L^-1^), which may be contributing to the abundance of hydrogenotrophic methanogens at D2 (see **Fig 3** and **Fig S1**). Acetate concentrations, as a proxy for the availability of organic acids produced in phase 2, were an order of magnitude higher at D2 vs D1 prior to sampling (i.e., 74.0 mg L^-1^ vs 2.3 mg L^-1^, respectively, as of January 2019, see **Fig S1**). These observations suggest that D2 is transitioning from phase 3 to phase 4 and frame D2 as a potential methane production hotspot whose contributions may be masked when examining bulk gas flaring data at the scale of the entire landfill cell.

#### Landfill cell E

Landfill cell E was originally classified as being in **phase 3 of rapid methanogenesis**. The low abundance and diversity of putative methanogens in cell E is at odds with the original classification based on geochemistry. Only seven putative methanogenic MAGs were recovered from landfill cell E, from the families *Methanotrichaceae*, also observed in cells A, B, C, and location D1; *Methanofollaceae*, also observed in D2; and the *Methanobacteriaceae* and *Methanospirillaceae* families, neither of which had previously been identified at this landfill (**Fig 4**). One MAG could only be classified to the *Methanobacteriales* order (see **Fig 3** and **File S1**).

Two MAGs from the family *Methanotrichaceae* displayed coverage values higher than the median and mean coverage observed at the whole-community level (i.e., 22.43 for STE_164 and 16.00 for STE_41 vs 5.85 for the median and 11.50 for the mean), whereas the remaining putative methanogen MAGs had lower coverages, ranging from 2.67 to 6.72 (**Fig 3**).

The methanogenic capabilities potentially supported in landfill cell E show a divide between acetoclastic and hydrogenotrophic methanogens. Acetoclastic methanogens were consistently more abundant than their hydrogenotrophic counterparts (**Fig 3**).

We expected cell E to be a favourable habitat for methanogens. The occurrence of acetoclastic and hydrogenotrophic methanogens in cell E aligns with our initial prediction that a limited number of substrates capable of supporting methane production would be available at this stage in the landfill’s lifecycle; however, the observed low abundance and diversity of methanogens was unexpected. The methanogenic community in cell E resembles those associated with waste that is 11 to 19 years older (i.e., cells A, B, and C) (**Fig 3**). From a microbiological perspective, these observations suggest cell E has reached the end of phase 3 and will soon enter phase 4 of slow methanogenesis.

The revised classification of cell E is difficult to reconcile with the consistently high methane production associated with this part of the landfill. Potential explanations are threefold: First, the abundance of putative methanogens may be decoupled from their metabolic activity. However, we would expect increased rates of methanogenesis to translate into more biomass and/or genomes given that methanogenesis is energy-conserving. Second, the sharp declines in inorganic carbon that occurred just prior to sampling (see **Fig S1**) may have caused a decline in acetoclastic and hydrogenotrophic methanogens using inorganic carbon for methane production. Finally, our characterization of the methanogen community from leachate at a more local scale may not align with gas data obtained for the entire landfill cell. Given that cell E is the only cell that experienced extreme fluctuations in substrates that could support methane production at the time of sampling, we’re inclined to attribute observed disparities to the availability of bicarbonate, though this mechanism needs to be formally tested in the future.

#### Landfill cell F

Landfill cell F was originally classified as **transitioning from phase 2 of anaerobic acid production to phase 3 of rapid methanogenesis**. In contrast to the samples obtained from both locations in cell D, the abundance, diversity, and community structure of putative methanogens was more consistent between both locations sampled from cell F despite their two-year age difference. 20 putative methanogen MAGs were recovered from location F1 and 17 putative methanogen MAGs were recovered from location F2. The families detected at landfill cell F (i.e., *Methanobacteriaceae, Methanocorpusculaceae, Methanocullaceae, Methanofollaceae, Methanomethylophilaceae, Methanomicrobiaceae, Methanoregulaceae, Methanosarcinaceae*, and *Methanotrichaceae*) were all detected in the landfill cells discussed previously (**Fig 3**). There was considerable overlap in the families detected in F1 and F2 aside from location F1 harbouring MAGs from the *Methanofollaceae* and *Methanomicrobiaceae*, families not detected at F2 (**Fig 3**).

Putative methanogen MAGs from location F1 displayed high relative abundance, with most displaying coverage values higher than the median and means obtained at the whole-community level (**Fig 3**). Most MAGs from the F2 methanogenic community displayed coverage values comparable to the median and mean coverage values obtained at the whole-community level suggesting they comprised a lower fraction of the total microbial community compared to those from F1 (**Fig 3**). At both F1 and F2, MAGs from the *Methanocullaceae* family were the most abundant putative methanogens, suggesting this family is well-suited to landfill habitats primed for rapid methane production (**Fig 3**).

The putative methanogenic communities from locations F1 and F2 displayed similar pathways that could support methanogenesis, with strict hydrogenotrophic and acetoclastic pathways represented in multiple families. All MAGs from the family *Methanosarcinaceae* could be classified as broad substrate methanogens possessing near complete pathways for acetoclastic and hydrogenotrophic methanogenesis alongside genes required to convert methyl-bearing substrates to methane (i.e., STF1_134, STF1_80, STF1_17, STF1_12, STF2_199, STF2_28, STF2_135) (**Fig 3**). The predicted capacity for multiple methanogenic pathways in members of the *Methanosarcinaceae* in cells F and D suggest members of this lineage are generalists capable of accessing a variety of substrates to support methane production over the course of a landfill’s lifecycle. The absence of the *Methanosarcinaceae* in cells A, B, and C also suggests that generalists from this family are outcompeted by more specialized methanogens as MSW ages (**Fig 4**).

The sole MAG classified to the order *Methanofastidiosales* (i.e., STF1_149) carried a homolog for the *mtsA* gene coding a methyltransferase specific to methylthiol-bearing compounds (see **Supplemental Results**). Members of this order carry out methanogenesis in a fastidious manner via the reduction of methylated thiols and we predict the same metabolism here (see **Supplemental Results**) (35).

We note MAGs from the abundant *Methanocullaceae* family were mixed as to their classification as hydrogenotrophic or acetoclastic methanogens (see **Supplemental Results**). These predictions merit further investigation as previous work on members of the *Methanoculleus* genus, the only genus of *Methanocullaceae* recovered from the landfill, suggested they are incapable of acetoclastic methanogenesis (see **File S1**) (36, 37).

The abundance and diversity of methanogens occurring in cell F aligns with our original classification of this landfill as transitioning from phase 2 of anaerobic acid production to phase 3 of rapid methanogenesis. Putative methanogens were more abundant than most other taxa detected in these parts of the landfill, likely increasing in abundance as conditions conducive to fermentation give way to those better suited to methanogenesis. Despite the peaks in organic acids being considerably lower in cell F compared to older landfill cells, the presence of varied substrates may support rapid methanogenesis due to limited competition for carbon substrates among methanogens (**Fig S1**). We posit that generalists and specialists alike have ample resources to support methanogenesis and contribute to high rates of methane production in cell F.

#### Methanogen families occurring in landfills

To place the methanogenic taxonomic diversity found at our study site in context, we generated a compilation of presence/absence data for methanogenic taxa identified in landfills in the past 20 years (**Fig 4**). Our study site harbours some of the most diverse communities of methanogens reported to date (i.e., 11 families detected). Only a 16S rRNA amplicon sequencing survey conducted across six distinct landfills in China showed higher diversity (i.e., 14 families detected, see **Fig 4**), which displayed considerable overlap in the families observed at our site (19). The remaining studies compiled, many of which also analyzed more than one landfill, detected lower taxonomic diversity regardless of the method employed (**Fig 4**).

Members from the *Methanosarcinaceae* are the most frequently detected in landfills, occurring in 16 of the 20 studies included (**Fig 4**). Members of the *Methanosarcinaceae* are widespread in terrestrial, aquatic, and animal-associated habitats due to their ability to use multiple substrates to support methanogenesis (38, 39). This generalist strategy likely contributes to their occurrence in landfills covering a range of ages and geographic locations, and frames members of the *Methanosarcinaceae* as key players in the landfill methane cycle (see **File S4**).

After the *Methanosarcinaceae*, the *Methanotrichaceae* (previously referred to as the *Methanosaetaceae* in the literature) (14/20 studies), *Methanocullaceae* (13/20 studies), and *Methanobacteriaceae* (12/20 studies) families were the next most frequently detected in landfills (**Fig 4**). Based on our own metabolic models, these families largely contain acetoclastic methanogens although there is conflicting information as to whether *Methanobacteriaceae* and *Methanocullaceae* are hydrogenotrophic methanogens or acetoclastic methanogens in the literature (38, 40–44). These families were repeatedly detected in newer waste but also waste that was over 20 years old, including in our data, suggesting they are important contributors to long-term methane production in landfills (see **File S4**).

The persistence of families such as the *Methanotrichaceae* and *Methanocullaceae* in older waste could be attributable to their increased tolerance to oxidative stress (45), an important adaptation to heterogeneous landfill habitats subject to large fluctuations in redox potential. Indeed, many of the methanogen families detected in landfills fall within orders forming a distinct clade of methanogens whose genomes are enriched in oxidative stress tolerance genes (e.g., *Methanocorpusculaceae, Methanofollaceae, Methanomicrobiaceae, Methanoregulaceae, Methanosarcinaceae, Methanosphaerulaceae, Methanospirillaceae*) (45) (**Fig 4**).

Families containing more metabolically restricted methanogens were less frequently detected in landfills. Members of the *Methanofollaceae* and *Methanocorpusculaceae*, considered to be strictly hydrogenotrophic methanogens, were detected in 8 and 4 of 20 studies, respectively (46–48) (**Fig 4**). Strictly methylotrophic methanogens from the family *Methanomethylophilaceae* were detected in 4 of 20 studies and methanogens from the family *Methanofastidiosaceae*, known for their use of methylthiols in methane production, were only detected in one study, suggesting landfill habitats are generally not conducive to supporting this specific methanogenic lifestyle (**Fig 4**) (34, 35, 49).

These observations do not preclude metabolically restricted methanogens from being important contributors to methane production in landfills. Strictly hydrogenotrophic methanogens were some of the most abundant MAGs recovered from location D2 in our study and likely play an important role in producing methane as organic substrates are depleted. Likewise, multiple MAGs of strictly methylotrophic methanogens from the *Methanomethylophilaceae* family were detected in cell F with coverage values above the median for the whole community, suggesting they are active contributors to methane production earlier in the landfill lifecycle when labile organic substrates are more likely to be available. The contributions of methylthiol-using methanogens to the methane cycle remains poorly characterized although the detection of members of the *Methanofastidiosales* in two different landfill studies suggest they also contribute to methane production in MSW.

### Methanotrophy

Our initial survey of MAGs containing the *pmoA* and/or *mmoX* genes identified **31 MAGs** as putative aerobic methanotrophs, with the addition of the **2 ANME MAGs** from D1 as putative anerobic methanotrophs. Within the aerobic methanotrophic MAGs, 15 encoded only *pmoA*, 5 encoded only *mmoX*, and 11 carried genes for both pMMO and sMMO complexes (**Fig 5**). Almost all putative methanotroph MAGs were found in parts of the landfill where oxygen was detected or suspected to be intruding (i.e., cells A, B, C, and location D1; see **Fig 2** and **Fig 5**).

**Figure 5:**
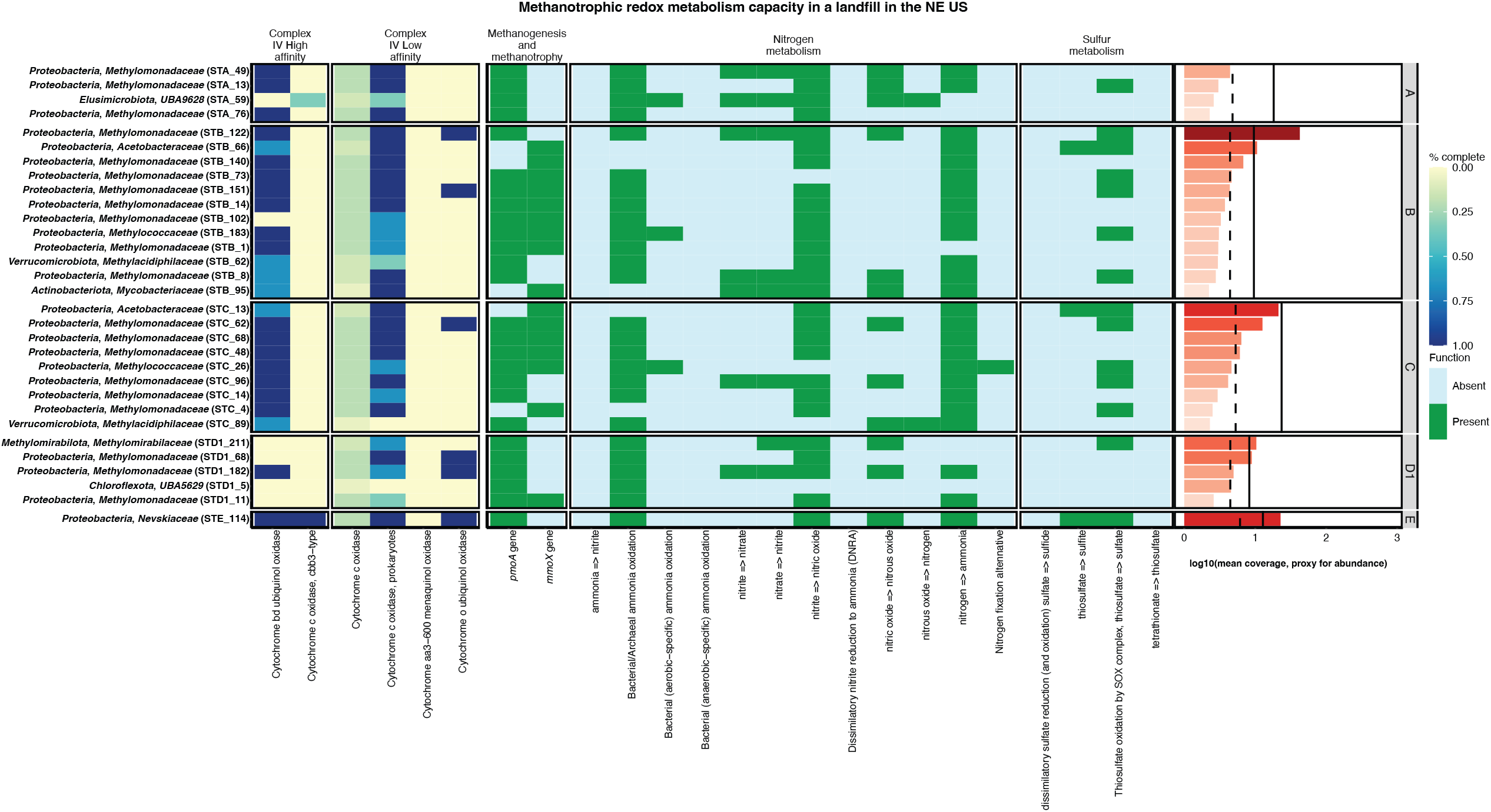
Metabolic heatmap and coverage data for putative methanotrophic MAGs. Shown here is the completion of high and low affinity complex IV machinery involved in reducing oxygen alongside the presence/absence of genes involved in methane metabolism, nitrogen metabolism, and sulphur metabolism. No putative methanotroph MAGs were recovered from location D2, F1, and F2, and as such no panels for those sampling locations are shown. Coverage values denote the mean coverage of all scaffolds that fell within a putative methanotrophic MAG under a log_10_ transformation. Solid black vertical lines denote the mean coverage calculated for MAGs at the whole-community level and dashed black lines denote the median coverage calculated for MAGs at the whole-community level.

Putative methanotroph MAGs spanned all three phyla with known methanotrophic representatives: the *Proteobacteria, Verrucomicrobiota*, and *Methylomirabilota* (50, 51). Of these, 25 MAGs belonged to families associated with aerobic or intra-aerobic methanotrophy, including the *Methylomonadaceae* (20), *Methylococcaceae* (2), *Methylacidiphilaceae* (2), and *Methylomirabilaceae* (1). Six MAGs were recovered from lineages whose capacity for methanotrophy remains poorly characterized or untested. These MAGs were taxonomically classified within the *Proteobacteria* families *Acetobacteraceae* (i.e., STB_66 and STC_13) and *Nevskiaceae* (STE_114), in addition to the phyla *Elusimicrobiota* (STA_59), *Actinobacteriota* (STB_95), and *Chloroflexota* (STD1_5) (**Fig 5**).

#### Methanotrophic community structure

In landfill cell A, four putative methanotroph MAGs were detected. Three MAGs belonged to the family *Methylomonadaceae* (i.e., STA_49, STA_13, and STA_76) and one belonged to the unnamed family UBA9628 from the phylum *Elusimicrobiota* (STA_59) (**Fig 5**). All MAGs displayed similarly low abundance with coverage values ranging from 2.29 to 4.42 (see **Fig 5** and **File S1**). The coverage values for putative methanotrophs were slightly lower compared to putative methanogens, which ranged in coverage from 3.32 to 6.88 (see **Fig 3, Fig 5**, and **File S1**), suggesting methane cycling is balanced but not a dominant process in this landfill cell.

Putative methanotroph MAGs were more abundant and diverse in landfill cell B compared to cell A, with 12 MAGs recovered from cell B. Ten of these MAGs were classified to families with known methanotrophic activities [i.e., *Methylomonadaceae* (8), *Methylococcaceae* (1), and *Methylacidiphilaceae* (1)] (51). The remaining two MAGs were classified to the *Acetobacteraceae* from the phylum *Proteobacteria* (STB_66) and the family *Mycobacteriaceae* from the phylum *Actinobacteriota* (STB_95) (**Fig 5**). MAG STB_122 from the *Methylomonadaceae* was the most abundant among the putative methanotrophs with a coverage value of 42.50 (see **Fig 5** and **File S1**). The remaining putative methanotrophic MAGs had coverage values ranging from 2.23 and 10.62, similar to the range reported for the methanogenic community (4.88 to 8.75) and most other microbial taxa in cell B (see **Fig 3, Fig 5, Fig S2**, and **File S1**).

The putative methanotrophic community from cell C displayed a similar pattern to cell B with respect to abundance and diversity. Nine putative methanotrophic MAGs were recovered from cell C. Eight of these MAGs were classified to families with known methanotrophic activities [i.e., *Methylomonadaceae* (6), *Methylococcaceae* (1), and *Methylacidiphilaceae* (1)] (51) and one MAG was classified to the family *Acetobacteraceae* (STC_13) (**Fig 5**). MAG STC_13 from the *Acetobacteraceae* was the most abundant putative methanotroph with a coverage value of 21.27 (see **Fig 5** and **File S1**). The remaining putative methanotrophs had coverage values ranging from 2.30 to 12.66, similar to the methanogenic community (i.e., 2.60 to 13.16) (see **Fig 3, Fig 5**, and **File S1**).

The detection of two *Methylacidiphilaceae* populations in cells B and C is of note because outside of a single amplicon sequencing study that detected this family in a corroding sewer pipe (52), the *Methylacidiphilaceae* have been exclusively characterized in acidic geothermal habitats [compiled in review by (53)]. Phylogenomic analyses shows that MAGs STB_62 and STC_89 are distinct from each other and clustered within the *Methylacidimicrobium* genus, which is considered to be a “mesophilic” clade within the *Methylacidiphilaceae* (53–56) (**Fig S3**). Metabolic reconstructions identified the presence of urease genes in the two landfill *Methylacidiphilaceae* MAGs that were absent in other *Methylacidiphilaceae* genomes (see **File S5** and **Supplementary Results**). The capacity to hydrolyze urea may represent an adaptation to landfills where the hydrolysis of urea could fulfill the dual role of providing a nitrogen source and inorganic carbon to support biomass production through the Calvin-Benson-Bassham cycle, as seen in geothermal representatives of the *Methylacidiphilaceae* (54, 57) (see **File S6**). Our observations challenge the view that members of the *Methylacidiphilaceae* are exclusively acidophiles and indicate this family exists in temperate circumneutral habitats where methane is available.

Sampling location D1 displayed lower abundance and diversity of putative methanotroph MAGs compared to cells B and C, with three MAGs classified to the *Methylomonadaceae*, one classified to the unnamed family UBA5629 in the phylum *Chloroflexota* (STD1_5), one classified to the family *Methylomirabilaceae* (STD1_211), and the two ANME *Methanoperedenaceae* MAGs (see **Fig 3** and **Fig 5**). MAG STD1_6 from the *Methanoperedenaceae* was the most abundant putative methanotroph MAG with a coverage value of 19.74 (see **Fig 3, Fig 5**, and **File S1**). The remaining MAGs for putative methanotrophs displayed coverages ranging from 2.60 to 10.37, similar to the range reported for methanogens (i.e., 3.05 to 6.73; see **Fig 3, Fig 5**, and **File S1**). No methanotrophic lineages were identified in location D2.

Finally, landfill cell E harboured a single putative methanotroph MAG from the family *Nevskiaceae* (STE_114) (see **Fig 5**). This MAG had a coverage value of 22.72, which was very close to the most abundant putative methanogenic MAG STE_164 from the family *Methanotrichaceae* (22.43; see **Fig 3, Fig 5**, and **File S1**). To the best of our knowledge, members of the *Nevskiaceae* family have yet to be tested for their ability to carry out methanotrophy through the pMMO pathway (see **Fig 5**). No methanotrophic lineages were identified from cell F.

#### Expanding the breadth of methanotrophic niches in landfills

The occurrence of the *Methylomirabilaceae*, the only family thought to support intra-aerobic methane oxidation, alongside the *Methanoperedenaceae*, suggests landfills (here location D1) are conducive to methanotrophic lifestyles that do not rely on exogenous oxygen. Although previous work has demonstrated the anaerobic oxidation of methane (AOM) in landfill cover soil microcosms and anoxic landfill sites, the contributions of anaerobes to methane oxidation remain understudied compared to aerobes in cover soils (58, 59). We identified *Methanoperedenaceae* and *Methylomirabilaceae* populations that can potentially contribute to AOM in the landfill and metabolic adaptations in putative aerobic methanotrophs that may support methane oxidation via the pMMO and sMMO pathways coupled to anaerobic metabolism.

Phylogenomic analyses of the *Methanoperedenaceae* showed that the two MAGs recovered from the landfill were distinct from each other, clustering with unnamed *Methanoperedens* species sequenced from contaminated groundwater and bioreactor enrichments (**Fig S4**). MAG STD1_6, which had the highest coverage among all *mcr*-containing MAGs recovered from location D1, may represent a member of the *Methanoperedenaceae* with distinct adaptations to contaminated aquatic habitats that merit further investigation (**Fig 3**).

Previous work has shown that members of the *Methanoperedenaceae* can oxidize methane via reverse methanogenesis coupled to the reduction of metals, oxidized nitrogen, and oxidized sulphur with the help of sulphate reducing partners (60–63). The *Methanoperedenaceae* MAGs recovered from the landfill displayed differing redox metabolisms that could potentially be coupled to AOM. MAG STD1_6 lacked the potential to reduce oxidized forms of nitrogen and sulphur, like the unnamed species *Methanoperedens* sp. 902386115, which is curious given that *Methanoperedens* sp. 902386115 is more closely related to MAG STD1_19 (see **Fig S4** and **File S6**). MAG STD1_19 demonstrated the potential to reduce nitric oxide to nitrous oxide, a trait observed in 7 of the 13 *Methanoperedenaceae* genomes analyzed (see **File S6**). MAG STD1_19 lacked the genes required for other steps in denitrification, suggesting it would require a metabolic handoff to acquire nitric oxide as a metabolic substrate to support AOM (see **File S6**). Alternatively, it’s possible these organisms use extracellular electron transfer (evidence supporting this physiological mechanism is discussed further in the **Supplemental Results**)

The *Methylomirabilaceae* MAG STD1_211 was most closely related to *Methylomirabilis limnetica*, the only genome for this lineage sequenced from a freshwater habitat to date (**Fig S5**). These two genomes cluster with *Methylomirabilis* sp.002634395 as a sister clade to *Methylomirabilis* genomes associated with metal-amended bioreactors and ditch sediments (**Fig S5**).

MAG STD1_211 encodes the capacity to reduce nitrate to nitrous oxide in line with all other *Methylomirabilaceae* genomes analyzed (see **File S6**). This observation supports that the *Methylomirabilaceae* could participate in the metabolic handoff required by the *Methanoperedenaceae* populations found at similar abundance at the same location. The potential for syntrophism between the *Methylomirabilaceae* and *Methanoperedenaceae* populations is supported by previous work where these families co-occur in systems where AOM occurred in the presence of oxidized nitrogen (64, 65).

The potential to couple methane oxidation to the reduction of oxidized nitrogen species was not limited to MAGs from the *Methanoperedenaceae* and *Methylomirabilaceae* families. The majority of putative methanotrophic MAGs in our dataset (27/31) demonstrated the capacity to reduce nitrite to nitric oxide and a smaller proportion could reduce nitrate to nitrous oxide (8/31) (**Fig 5**). These traits spanned multiple families (e.g., *Acetobacteraceae, Methylacidiphilaceae, Methylococcaceae, Methylomirabilaceae, Methylomonadaceae, Mycobacteriaceae*, and *Nevskiaceae*) (**Fig 5**). Although the methanotrophic capacity of members from the *Acetobacteraceae* and *Nevskiaceae* has yet to be experimentally confirmed, these MAGs (i.e., STB_66, STC_13, and STE_114) displayed the capacity to reduce thiosulfate as a potential electron acceptor that could be coupled to methane oxidation (**Fig 5**). The majority of putative methanotrophic MAGs (24/31) also displayed high completion for the high affinity complex IV capable of scavenging nanomolar levels of oxygen (66–68), and a similar proportion (26/31) displayed high completion for the low affinity complex IV (**Fig 5**). These observations support that pMMO and sMMO-bearing methanotrophs in leachate have versatile redox metabolisms that can help them survive in landfill habitats separated from the atmosphere.

These adaptations could allow methanotrophs to generate energy in the absence of oxygen as a terminal electron acceptor while also allowing methanotrophs to reserve trace amounts of oxygen to support methane oxidation via the pMMO and/or sMMO pathways. Similar explanations have been put forward for observations where members of the *Methylomonadaceae* were abundant in anoxic aquatic habitats associated with high rates of methane oxidation separate from oxic-anoxic interfaces where aerobic methane oxidation is typically thought to occur (69, 70). Notably, members of the *Methylomonadaceae* have also been detected in studies of landfill cover soils and bioreactors inoculated with leachate subject to oxygen gradients, providing a good proxy for the range of redox conditions considered in our study (9, 23, 25). These observations and the widespread detection of *Methylomonadaceae* in leachate frame members of this family as key players capable of limiting methane emissions over a wide range of redox potentials in landfills.

#### Phylogenomic analyses of families with no known capacity for methanotrophy

The detection of putative methanotrophs in the *Nevskiaceae, Acetobacteraceae*, which are families lacking *in vivo* evidence of methanotrophy, and the *Mycobacteriaceae*, which displayed conflicting physiological evidence surrounding the capacity for methanotrophy until very recently (71), prompted us to examine whether the methanotrophic potential observed for these genomes was unique to the landfill-derived populations or more widespread in each lineage. Phylogenomic analyses revealed that multiple genomes spread throughout each family’s tree carried the genes coding for complete pMMO and/or sMMO complexes, suggesting the capacity for methanotrophy has been acquired on several occasion within each lineage (See **Supplemental Results** for detailed descriptions; **Fig S6, S7, S8**). Many genomes also displayed near-complete sMMO complexes but consistently lacked the *mmoZ* gene, possibly due to the *mmoZ* gene being prone to divergence as seen in the genomes of known methanotrophs (72) (see **Fig S6, S7**, and **S8**). Outside of the *Mycobacterium holsaticum* genome associated with human sputum, almost all putative methanotrophs identified across the three lineages occurred in aquatic, terrestrial, and sediment habitats where methane oxidation occurs (27, 51).

These observations suggest that many of these lineages are potentially overlooked as methane oxidizers in these habitats. We highlight the *Mycobacteriaceae* as a case study in the **Supplementary Results** to place our phylogenomic analyses in the context of the recent discovery of methanotrophy in an isolate from this family, which clarifies the previous conflicting reports concerning methane oxidation in representatives of this family (71). In the future, it will be important to take advantage of the fact that many of these strains, in the *Mycobacteriaceae* but also from the other lineages profiled here, exist in culture collections, providing an opportunity to validate their predicted methane oxidation capacity.

## Conclusions

This study provides an unprecedented historical perspective on biogeochemical succession and the resultant shifts in methane cycling guilds over decades of MSW ageing. Our findings show that geochemical monitoring captures the major processes happening in the landfill but lacks nuance from microbiological information essential to predicting methane’s fate in a landfill.

Our metagenomic analyses showed that newer landfill habitats support more abundant, diverse, and metabolically versatile communities of putative methanogens compared to older habitats. Methanogen community structure in ageing MSW seems to be controlled by the variety and availability of substrates that can support methane production, as well as oxygen infiltration inhibiting methanogenesis. Methanotrophs had more restricted distribution and diversity. When present, methanotrophs tended to occur at similar or slightly higher abundance than methanogens. We observed the capacity for methanotrophy across a broad range of metabolisms that likely reflect the steep redox gradients encountered in landfills. The widespread adaptations observed in central redox metabolisms suggests that methanotrophy, even via oxygen-requiring pathways, is important to consider in anoxic landfill habitats.

One of the most exciting findings that emerged from our metagenomic analyses is that pathways and microbial taxa involved in anaerobic oxidation of methane are more diverse than previously described. The abundance of anaerobic methane oxidizers at location D1 in this study raises important questions as to what variables result in a habitat favouring anaerobic vs aerobic pathways for methane oxidation. Identifying these variables will be crucial for developing biostimulation strategies that allow landfills to function as giant anaerobic methane oxidizing bioreactors once methane production has dropped below sustainable bio-energy generation levels. Our work indicates that it is important to dig below cover soils, into the anoxic habitats that dominate landfills, to expand the current concept of the niches and diversity of microbial taxa that contribute to methane oxidation in MSW. Although physiological experiments are required to confirm the methane oxidation capacity in the three different families containing novel putative methanotrophs, this discovery reinforces how the unique microbial communities in landfills can help us better understand biogeochemical cycles across different habitats.

Expanding the known diversity of methane cycling microorganisms is crucial for improving biogeochemical models that can be used to manage methane emissions in landfills. Such models could increase the effectiveness of waste diversion programs, identify substrate amendments for optimal methane productions and recovery for landfills, or limit methane emissions based on waste composition for landfills with higher emission profiles. Our study advocates for emphasizing the biological dimension of landfill lifecycle models so that these tools are not only used for monitoring but actively mitigating the negative environmental impacts of MSW degradation.

## METHODS

### Leachate collection and geochemical analyses

Leachate sampling involved purging wells of standing liquid prior to using a peristaltic pump to recover 1 L of leachate in sterile Nalgene bottles. Samples were stored on ice prior to filtration through a 0.22 µm pore size sterivex filter (Millipore Sigma, Burlington, MA). Leachate was filtered until the filter clogged, after which sterile air was forced through the filter to remove residual leachate. Two to three sterivex filters were used for each leachate sample to ensure sufficient biomass recovery for downstream analyses. Leachate filtration was performed same day as sampling and filters were stored at −80ºC until processed for DNA.

Monitoring records for leachate geochemistry were kindly provided by site owners dating back to 1983 (i.e., 36 years of records). Additional gas flaring data associated with each landfill cell were also provided for the years 2018 and 2019 to capture historical seasonal variation in gas production over a time frame relevant to the sampling expedition. Select variables, namely BOD, COD, pH, redox potential (ORP), the concentrations of organic acids (i.e., acetic acid, butyric acid, isobutyric acid, propionic acid, and valeric acid), bicarbonate (i.e., a proxy for dissolved inorganic carbon), volume of gas flared, and the composition of the gas flared were used to classify each landfill cell to a specific phase (i.e., phase 1 to 5) in the landfill conceptual model.

### DNA extraction, metagenomic sequencing, and genome assembly

DNA was extracted from biomass on filters using the PowerSoil DNA Isolation Kit (Qiagen) using the manufacturer’s protocol with the exception that diced filters were added to the bead tube in place of soil. Extracted DNA was evaluated for quality using the NanoDrop 1000 (Thermo Scientific, Waltham, MA) and quantified using the Qubit fluorometric method (Thermo Scientific, Waltham, MA) following the manufacturer’s protocol.

Extracted DNA for all samples was sent to The Center for Applied Genomics (Toronto, Canada) for shotgun metagenome sequencing using an Illumina HiSeq platform with paired 2 × 150 bp reads (Illumina, San Diego, CA). Metagenomic reads were quality trimmed using bbduk in the BBTools suite (https://sourceforge.net/projects/bbmap/) and Sickle v1.33 (73) and reads were subsequently assembled into scaffolds using SPAdes3 v.3.15.5 (74) using-meta and kmers set to 33,55,77,99, and 127. Only scaffolds greater than or equal to 2.5 kbp in length were further analyzed. Metagenomic reads were mapped separately to each curated scaffold assembly using Bowtie2 v 2.3.4.1 (75).

Scaffolds from a single metagenome were binned using three binning algorithms: CONCOCT v0.4.0, MaxBin2 v2.2.6, and MetaBAT2 v2.12.1 (76–78). The resulting bins were dereplicated for each landfill leachate sample and scored in a consensus-based manner using DAS Tool v1.1.1 (79). To assess bin quality, DAS Tool-processed bins were used as input in CheckM v1.0.13 (80), which yielded 1,881 metagenome-assembled-genomes (MAGs) with >70% completion and <10% contamination retained for further analyses (**File S3**).

Mean coverage for a MAG was used as a proxy to compare the abundance of different populations across the landfill. Mean coverage was calculated by taking the mean of the coverage values reported for unique scaffold identifiers within a given bin. The distribution of the mean coverage values obtained for each MAG is summarized in **Figure S2** and the individual mean coverage values associated with each MAG can be found in **File S1**.

### Genome taxonomy, annotation, and metabolic summaries

Taxonomy was assigned to MAGs using the Genome Taxonomy Database Toolkit application (GTDB-tk) r89 available on DOE-KBase (81). MAGs were annotated using the Distilled and Refined Annotation of Metabolism (DRAM) tool v1.0 with default parameters (82) but omitting the use of the KEGG and UniRef90 databases for initial annotation of all 1,881 MAGs. In specific cases where <20 genomes from a specific lineage needed to be characterized in additional detail, the UniRef90 database was applied to test whether it improved annotations for key pathways (see **File S5, File S6**, and **File S7**). DRAM generates an annotation file that was used alongside the product file and taxonomy data to identify putative methanogens and methanotrophs through additional data manipulation in R v4.1.2 (see **File S1, File S2**, associated R code used to produce all figures can be found at https://github.com/carleton-envbiotech/Methane_metagenomics).

### Methane cycling microbial community analyses

Detailed guidelines on how DRAM’s output and GTDB taxonomic classification were used to characterize putative methanogenic and methanotrophic MAGs have been provided in **Supplemental Methods**. Briefly, we used DRAM’s product file to identify putative methanogen MAGs by verifying whether MAGs possessed the *mcrA* gene or a nearly complete pathway for hydrogenotrophic methanogenesis (see **File S1** and **Supplemental Methods**). MAGs classified to methanogenic lineages but lacking the *mcrA* gene had their annotations verified manually for other genes from the *mcr* operon prior to further analyses (see **File S2** and **Supplemental Methods**). Putative methanogen MAGs were subsequently classified as strictly hydrogenotrophic, acetoclastic, strictly methylotrophic, or capable of using a broad array of substrates to support methane production using information contained in DRAM’s product file (see **File S1** and **Supplemental Methods**).

To facilitate comparisons to previous work, we compiled presence/absence data for methanogenic taxa reported in previously published landfill studies. We used a permissive definition for indicating a taxon was present, wherein measures of relative abundance > 0% or taxa identified using qualitative tools such as fluorescent *in situ* hybridization were taken as evidence of presence (see **Supplemental Methods**). We normalized the taxa reported in the literature to the GTDB taxonomy release 89 originally used to taxonomically classify MAGs from our landfill site and have provided the input file used for this metanalyses in **File S4**. The associated code used to normalize taxonomy and produce visualizations can be found at https://github.com/carleton-envbiotech/Methane_metagenomics.

MAGs for putative aerobic methanotrophs were identified based on the presence of the *pmoA* and/or *mmoX* genes coding for key subunits in the pMMO and sMMO enzyme complexes in DRAM’s product file (see **File S1**). MAGs taxonomically assigned to ANME were manually identified and reclassified as predicted methanotrophs. In cases where putative methanotrophs were taxonomically classified to families with no previous record of methanotrophy, annotations were manually verified to ensure multiple *pmo* and *mmo* genes occurred on scaffolds with minimum lengths of ~3000 bp prior to subsequent analyses (see **File S2**).

### Phylogenomic analyses

All phylogenomic analyses were conducted with GToTree v1.5.38 (83), which references the GTDB release 202 taxonomic identifiers (81, 84) when retrieving publicly available genomes. GToTree was used to collect representative genomes within a lineage of interest and related lineages to build outgroups. GToTree automatically checks the quality of input genomes by quantifying representation of curated sets of single copy genes.

To visualize distribution of methane oxidation marker genes, Pfam identifiers for the pMMO and sMMO pathways were supplied to GToTree. The metadata file output by GToTree with *pmo* and *mmo* gene counts was subsequently used to establish strict criteria for identifying genomes as encoding the pMMO, sMMO, or pMMO and sMMO pathways.

Categories were overlaid onto phylogenetic trees in R v4.1.2 using the packages ‘ggtree’ v3.0.4 (85, 86) and ‘treeio’ v1.16.2 (87). Isolation sources of genomes were manually retrieved using the NCBI biosample number associated with each genome and overlaid onto trees when pertinent. Accession numbers from GToTree were supplied to the ‘bit’ v1.8.53 package to download genomes, which were included as input for metabolic models produced using DRAM. Examples of R notebooks containing the code used to analyze each lineage of interest are provided alongside the input data required to reproduce these analyses at https://github.com/carleton-envbiotech/Methane_metagenomics.

## Supporting information

Supplemental methods and results including Figures S1-S8 and Table S1

## Data Availability

Sequencing data associated with this project have been deposited to NCBI under BioProject PRJNA900590. The raw reads files are available on the SRA database, under Biosamples SAMN31696084 – SAMN31696092. The 1,892 MAGs have been deposited to the WGS database under accessions SAMN32731718 – SAMN32731810 (STA), SAMN32731811 – SAMN32731998 (STB), SAMN32733587 – SAMN32733720 (STC), SAMN32734194 – SAMN32734413 (STD1), SAMN32734415 – SAMN32734683 (STD2), SAMN32734737 – SAMN32734946 (STE), SAMN32737191 – SAMN32737484 (STF1), SAMN32737485 – SAMN32737723 (STF2). The input DRAM annotation and product data and accompanying bash and R code used for the analyses in this manuscript has been provided as Supplementary Files for download via https://github.com/carleton-envbiotech/Methane_metagenomics. Geochemical monitoring records for leachate have been kept confidential at the request of site managers.

## Acknowledgements and Funding Sources

We are sincerely grateful to the landfill site management and their contracted consulting company (anonymity by request) for site access, aid with sampling, and provision of detailed monitoring records. We thank Dr. Jennifer Biddle and her lab group for hosting our team for sample processing prior to shipment. Thanks also to Ms. Rebecca Co and Ms. Alexandra Sauk for help with sampling. This work was supported by an NSERC Discovery Grant (2016-03686) to LAH. LAH was supported by a Tier II Canada Research Chair. DSG was supported by an NSERC Banting Postdoctoral Fellowship. NAG was supported by an NSERC CGS-M, an Ontario Graduate Scholarship, an NSERC PGS-D, and a W.S. Rickert Graduate Student Fellowship from the University of Waterloo.

