## Supplemental methods and results including Figures S1-S8 and Table S1 for "Microbial methane cycling in a landfill on a decadal time scale"

### Supplemental Information:
### Microbial methane cycling in a landfill on a decadal time scale

Daniel S. Grégoire, Nikhil A. George, Laura A. Hug

**Supplementary Methods**

- Extended rules for classifying genomes as putative methanogens
- Literature review method for compiling methanogenic taxa from landfill studies

**Supplementary Results**

- Quality screening of putative methanogen MAGs and the UBA148 family
- BLAST search for *mtaA/mtsA* in *Methanofastidiosales* metagenome-assembled genome
- Putative methanotrophs in the *Nevskiaceae*, *Acetobacteraceae*, and *Mycobacteriaceae*

**List of Supplementary Figures and Tables**

**Fig S1:** Organic acid concentrations through time for landfill cells from 1986 to 2019.

**Fig S2:** Density plots for mean coverage values calculated for metagenome-assembled-genomes.

**Fig S3:** Rooted tree for *Methylacidiphilaceae* genomes from the landfill and GTDB.

**Fig S4:** Rooted tree for *Methanoperedenaceae* genomes from the landfill and GTDB.

**Fig S5:** Rooted tree for *Methylomirabilaceae* genomes obtained from the landfill and GTDB.

**Fig S6:** Unrooted tree for genomes from the *Nevskiaceae* family identified as putative methanotrophs from the landfill and GTDB.

**Fig S7:** Unrooted tree for genomes from the *Acetobacteraceae* family identified as putative methanotrophs from the landfill and GTDB.

**Fig S8:** Unrooted tree for genomes from the *Mycobacteriaceae* family identified as putative methanotrophs from the landfill and GTDB.

**Table S1:** QA/QC summary for landfill metagenomes generated with BBMap.

**List of Supplementary Files (available for download via Github:** https://github.com/carleton-envbiotech/Methane_metagenomics)

File S1: Product file from DRAM appended with taxonomy and coverage data

File S2: Full annotation file (can be downloaded via https://www.dropbox.com/s/7g952zkox8y38rc/FileS2_DRAM_annotations_all_MAGs_names_redacted.tsv?dl=0 due to large size not supported by Github)

File S3: Completion contamination data

File S4: Raw compilation data for methanogenic taxa

File S5: Annotation file for the *Methylacidiphilaceae* genomes

File S6: Product files for *Methylacidiphilaceae, Methanoperedenaceae*, *Methylomirabilaceae*

File S7: Annotation file for *Methanoperedenaceae* genomes

#### Supplementary Methods

##### Extended rules for classifying genomes as methanogens and methanotrophs

DRAM's product file indicates whether genomes carry the *mcrA* gene essential to methane production, the level of completion for the hydrogenotrophic methanogenesis pathway, and whether genes required for using acetate, methanol, and amine-bearing molecules as substrates for methanogenesis are present in each genome. Our initial data filtration involved identifying MAGs with the *mcrA* gene **or** a 75 % complete pathway for hydrogenotrophic methanogenesis (equivalent to 6/8 steps being present in the "Methanogenesis, CO2 => methane" pathway output by DRAM) (see **File S1**). After this step, we used the data from DRAM's product and annotation files (**File S1** and **File S2,** respectively) and the taxonomic classification obtained from GTDB to categorize the metabolic strategies used by putative methanogens. We used the definitions and mechanisms summarized in an authoritative review on methanogens as the basis for these rules [(1) and references therein].

1. *Strictly hydrogenotrophic methanogenesis*: Strictly hydrogenotrophic methanogens can only produce methane by reducing carbon dioxide using hydrogen as an electron donor. MAGs identified as strictly hydrogenotrophic methanogens must have the *mcrA* gene and/or >75% completion of the hydrogenotrophic methanogenesis pathway. MAGs that lacked the *mcrA* gene but displayed high completion for the hydrogenotrophic pathway were further investigated to verify they were taxonomically classified to lineages with known methanogens and whether additional genes from the *mcr* operon were present. MAGs were included in analyses if they harboured any of the *mcrBCDG* genes found in the *mcr* operon. MAGs classified as strictly hydrogenotrophic methanogens also needed to lack the carbon monoxide dehydrogenase/acetyl-CoA synthase complex (CODH/ACS), denoted as "Acetyl-CoA pathway, $CO_2$ => Acetyl-CoA" in DRAM's output. The completion of this pathway is determined based on the presence of genes catalyzing reversible redox transformations between carbon monoxide and carbon dioxide, and methyl group transfers between the coenzyme M precursor tetrahydromethanopterin ($H_4MPT$) and acetyl-CoA (2, 3). This enzyme complex is a hallmark of acetoclastic methanogenesis but is also thought to support autotrophic carbon fixation in archaea bearing near complete hydrogenotrophic methanogenesis pathways that lack the *mcr* operon (3).

2. *Acetoclastic methanogenesis*: Acetoclastic methanogens convert acetate to acetyl-CoA, which subsequently undergoes dismutation to produce carbon dioxide and a methyl group. The carbon dioxide can be further converted to methane using the hydrogenotrophic pathway whereas the methyl group supplied from acetyl-CoA goes towards forming the precursor to coenzyme M, $H_4MPT\text{-}CH_3$. MAGs identified as putative acetoclastic methanogens needed to have the *mcrA* gene and/or genes coding for enzymes capable of converting acetate to acetyl-CoA. The potential to convert acetate to acetyl-CoA was evaluated based on the presence of genes coding for the acetyl-CoA synthetase alone (labelled as "Acetate pt 1" in DRAM's output), or the acetate kinase and acetyltransferase together (labelled as "Acetate pt 2 and 3", respectively). MAGs identified as putative acetoclastic methanogens also required a >50% complete CODH/ACS pathway. Given that CODH can further oxidize carbon monoxide to carbon dioxide, MAGs identified as acetoclastic methanogens also required > 75% completion for the hydrogenotrophic methanogenesis pathway. The annotation of MAGs displaying high completion of the hydrogenotrophic pathway alongside biomarker genes to convert acetate to acetyl-CoA but lacking the *mcrA* gene were examined to determine whether other genes in the *mcr* operon were present as a condition to being included in the dataset.

*3. Strictly methylotrophic methanogenesis*: Strictly methylotrophic methanogens can only produce methane using methylated compounds. MAGs identified as putative strictly methylotrophic methanogens needed to have the *mcrA* gene present alongside biomarker genes associated with the methyltransferase enzymes specific to methanol, trimethylamine, dimethylamine, and/or methylamine output by DRAM. DRAM does not output the presence of methylthiol transferases in the product file by default. In cases where methylthiol transferases were suspected as a potential pathway to support methanogenesis, the presence/absence of the *mtsA* and *mtaA* genes was manually verified in the annotation file for select genomes (4). MAGs identified as strictly methylotrophic methanogens were required to lack the CODH/ACS complex and display low completion (<50%) of the hydrogenotrophic methanogenesis pathway so that only methyl-bearing substrates could potentially support methane production.

4. *Broad substrate methanogenesis*: This category encompasses MAGs bearing the *mcrA* gene and the potential to access an array of substrates including inorganic carbon, acetate, methanol, and amine-bearing molecules alongside a >75% complete hydrogenotrophic pathway to produce methane.

**Compilation of presence/absence data for methanogenic taxa from landfill studies**

To compare the occurrence of methanogenic and methanotrophic taxa in this study to previous research, a meta-analysis was conducted for a select number of studies examining landfill microbial communities. The main criterion for inclusion was that these studies examined microbial communities *in situ* in landfills. Studies that sampled landfills to inoculate enrichment cultures were also considered but data was only collected if the original environmental sample was sequenced, and only that original environmental sample was used in the meta-analysis. Exceptions were made for temporal studies that did not apply selective forces to enrich for specific microbial guilds, but monitored the succession of microbial community associated with solid waste or leachate under conditions that support waste degradation *in situ* (5–7).

Presence/absence was determined by first examining the data presented in figures and tables in the published versions of articles. In specific cases where articles cited accessible supporting information, these data were also incorporated into the analyses. A liberal approach was taken for determining presence/absence. Specific taxa reported in the articles were recorded as being present. For amplicon sequencing surveys, which comprised the bulk of the data reported, any relative abundance >0 % for a given taxa was deemed sufficient to indicate that this taxon was present. For studies that used patterns in restriction fragment length polymorphism or closest relative matches to identify the taxa present, the name of the closest relative was recorded to indicate a taxon was present. For metagenomic studies, the taxa names associated with metagenome-assembled-genomes or biomarker genes used to assess abundance were recorded as those taxa being present. In instances where specific microarrays or fluorescent *in situ* hybridization were employed, the species names reported by the authors in the articles were taken as evidence of those taxa being present.

From all compiled data, the deepest level of taxonomic classification presented was recorded (see **File S4**). Given that our study focussed on comparing microbial communities at the family level, studies that did not provide taxonomic classification to the family-level or deeper were discarded from further analyses. Our study used GTDB release 89 as the database for taxonomic classification (8) whereas the data compiled from the literature relied on a variety of databases for 16S rRNA genes over several years (e.g., Greengenes, SILVA, NCBI) (9–11). To ensure consistent naming between the taxa compiled from the literature and our study, species or genus names compiled from the literature were manually searched in GTDB and the

GTDB name was compared to the NCBI name for searches that provided hits. The naming history of the taxon was also manually verified, such that a list of rules was developed to link older species, genera, and family names to the naming convention for families in GTDB release 89 (e.g., a name commonly reported in the literature was the family *Methanosaetaceae*, which is now *Methanotrichaceae* in GTDB). Conversions identifying the current family naming convention based on species, genera, and family names originally reported have all been reported in the R code that was used to visualize this data with comments linking the first occurrence of this name to a specific GTDB release (available under the directory "Methanogen_metaanalyses" via https://github.com/carleton-envbiotech/Methane_metagenomics).

#### Supplementary Results

##### Quality screening of predicted methanogen MAGs and the UBA148 family

Four MAGs from the unnamed family UBA148 were initially predicted to be methanogens. Upon curation, all MAGs from UBA148 were removed from the putative methanogen dataset because they lacked annotated *mcr* genes (see MAGs STB_81, STC_9, STD1_54, STD1_76, and STD2_203 in **File S2**). All other MAGs that were taxonomically classified to methanogen families but lacked the *mcrA* gene carried genes coding for subcomponent A2 of the methyl coenzyme M reductase and/or the *mcrC* and *mcrD* genes and were kept in the final dataset (see **File S2**).

According to GTDB release 89, UBA148 is in the order *Methanocellales*, which contains the methanogenic *Methanocellaceae* family (8, 12). Although MAGs belonging to the family UBA148 lacked *mcr* genes, they displayed >75 % completion of the hydrogenotrophic methanogenesis pathways alongside complete CODH/ACS complexes, and in some cases, the ability to convert acetate to acetyl-CoA (see **File S1** and **S2**). We also observed that the *mtrA* and *mtrH* genes from the *mtrEDCBAGH* gene cluster coding for the tetrahydromethanopterin S-methyltransferase, which produces methyl coenzyme M (13, 14), were present in UBA148 MAGs (see MAGs STB_81, STC_9, STD1_54, STD1_76, and STD2_203 in **File S2**).

In many instances, the abundance of UBA148 MAGs was higher than putative methanogens, particularly in older MSW (see **File S1**). We speculate that this may be tied to the confirmed or suspected oxygen infiltration in older landfill cells (i.e., A, B, and C). In manually verifying annotations for UBA148 MAGs, we observed that genes coding antioxidant enzymes including superoxide reductase, peroxiredoxin, and rubrerythrin, were present alongside genes associated with redox buffer systems such as thioredoxin and rubredoxin (see MAGs STB81, STC_9, STD1_54, STD1_76 in **File S2**). The presence of similar genes has been noted as potential adaptations to tolerating oxidative stress in bona fide methanogens (15).

The major difference between members of family UBA148 and methanogenic lineages in this study is that UBA148 genomes lacked genes coding for the *mcr* complex, which houses an extremely oxygen sensitive Ni(I) cofactor (16, 17). We speculate that members of the UBA148 family may gain a competitive advantage through the elimination of the oxygen sensitive *mcr* complex to support an autotrophic lifestyle in habitats where oxygen infiltration would directly inhibit *mcr*-dependent methanogenesis. This hypothesis has yet to be tested but underscores the need to understand how methanogens tolerate oxidative stress in ageing landfills and how these adaptations control long term methane production.

#### **BLAST search for *mtaA/mtsA* in *Methanofastidiosales* genomes**

In classifying the methanogenic capabilities observed in putative methanogen MAGs, we noted DRAM's product file showed that MAG STF1_149 from the order *Methanofastidiosales* had no genes demonstrating the potential to access inorganic or organic substrates to support methane production despite having an annotated *mcrA* gene (**Fig 3**). The first genomic characterization of the *Methanofastiodosales* suggested that members of this order carry out methanogenesis in a fastidious manner via the reduction of methylated thiols thanks to the *mtsA* gene coding a methyltransferase specific to methylthiol-bearing compounds (4).

To determine whether MAG STF1_149 from the order *Methanofastidiosales* could potentially support methanogenesis using the reduction of methylthiols via the *mtsA* gene, we conducted an in-depth analysis on this genome and others from the order *Methanofastidiosales* retrieved from GTDB. To confirm that DRAM could annotate the *mtsA* gene, we searched the raw annotations for all MAGs obtained from the landfill. We confirmed that the *mtsA* gene was annotated in 3 MAGs (i.e., STD2_188, STF1_134, and STF2_199) (see **File S2**). Notably, DRAM failed to annotate the *mtsA* gene for MAG STF1_149 (see **File S2**). Although failure to annotate the *mtsA* gene in MAG STF1_149 could be attributable to the low completion of this genome (see **File S3**), the *mtsA* gene could not be annotated in any publicly available *Methanofastidiosales* genomes even when including the UniRef90 database to improve annotations (data not shown).

The *mtsA*-encoding MAGs, STD2_188, STF1_134, and STF2_199, were all taxonomically classified to the species *Methanosarcina* sp.002499445 within the order *Methanosarcinales* (see **File S1**). Furthermore, 2 KEGG IDs were presented side by side for these annotations: K16954 for the *mtsA* gene, and K14080 for the *mtaA* gene (see **File S2**). The *mtaA* gene is a methyltransferase that catalyzes the final step of methylotrophic methanogenesis and is thought to be specific to methanol and/or amine molecules, whereas *mtsA* is thought to be specific to methylated thiols (4, 18, 19). This dual annotation suggests that DRAM is unable to distinguish between these two homologous genes using sequences available through reference databases such as KEGG, Uniref90, and Pfam. When verifying the literature references associated with the KEGG entry K16954 denoting the *mtsA* gene, both references associated with this entry are studies on *Methanosarcina barkeri* (references accessed via https://www.genome.jp/dbget-bin/www_bget?ko:K16954) and the taxonomy associated with the genes that KEGG refers to are dominated by members of the *Methanosarcina* genus. These observations suggest that DRAM's capacity to annotate *mtaA*/*mtsA* genes and potential homologs relies on sequences that are divergent from the *mtaA*/*mtsA* sequence of members of the *Methanofastidiosales*.

To identify putative *mtaA*/*mtsA* homologs in the *Methanofastidiosales* MAG STF1_149, we retrieved the *mtsA* sequence from the *Methanofastidiosum methylophilus* genome and performed a BLAST search using MAG STF1_149 as the searchable database. This search indicated that MAG STF1_149 harboured a gene with similarity score of 67% to the *mtsA* sequence of *Methanofastidiosum methylophilus*, which was annotated as a uroporphyrinogen decarboxylase by DRAM (corresponding to Pfam entry PF01208) (see **File S2**). Notably, the same PFam entry applies to the *mtaA* gene, further supporting that DRAM lacks the ability to distinguish between *mtaA* and *mtsA*. The similarity score being close to 70% further suggests that this gene in MAG STF1_149 may carry out the same function as *mtaA/mtsA* but may have diverged due to a specialized methanogenic lifestyle optimized for methylthiol-bearing compounds.

As an additional test for database biases limiting our ability to annotate *mtaA/mtsA* homologs in *Methanofastidiosales* genomes, the putative *mtaA/mtsA* sequence detected in MAG STF1_149 was used as input for a tblastn search to identify the taxa with the best hit. The best hit for the hypothetical *mtsA* gene in MAG STF1_149 had a similarity score of 68% and was associated with a sequence from *Methanosarcina acetivorans* C2A. Notably, no sequences from the *Methanofastidiosales* order showed up in the alignment table, despite the similarity score being nearly identical to those obtained with a BLAST search of MAG STF1_149 using the *mtsA* sequence from *M. methylophilus* as a query sequence. Regardless of whether the gene of interest in MAG STF1_149 represents a homolog for *mtaA* or *mtsA*, this result demonstrates that DRAM may not have been able to effectively annotate divergent sequences of this gene because it is referencing sequences in KEGG and Pfam originally obtained from the *Methanosarcina* genus. The inability to detect the *mtsA* gene with tools such as DRAM also suggests that divergent pathways for methanogenesis that rely on methylthiols remain overlooked in the methane cycle and merit further investigation. We conclude that there is evidence, albeit inconclusive, that MAG STF1_149 can carry out methanogenesis via a divergent *mtsA* homolog given that all other machinery to access known substrates for methanogenesis are absent.

**Putative methanotrophs in the *Nevskiaceae, Acetobacteraceae*, and *Mycobacteriaceae***

In the *Nevskiaceae* family, four genomes had all genes required for the pMMO complex (i.e., MAG STE_114 from this study and its close relative CAU-1509 sp. 005047655, *Panacagrimonas perspica*, and *Polycyclovorans* sp.002706265) and *Solimonas aquatica*'s genome had the full gene repertoire for both the pMMO and sMMO complexes (**Fig S6**). In the *Acetobacteraceae* family, four genomes carried the genes required for the sMMO complex (i.e.,

MAGs STC_13 and STB_66 from this study, which are closely related to each other, *Rhodopila* sp. 903915725 and *Rhodopila* sp. 903851415; **Fig S7**). In the *Mycobacteriaceae* family, which has substantially more publicly available genomes than the other two families examined, two genomes carried the genes required for the pMMO complex (i.e., *Mycobacterium dioxanotrophicus* and *Smaragdicoccus niigatensis*), five genomes carried the genes required for the sMMO complex (i.e., MAG STB_95 from this study, *Mycobacterium moriokaense* A, *Mycobacterium* sp. 003719305, *Mycobacterium holsaticum*, and *Mycobacterium pulveris*), and four genomes carried the genes required for both pMMO and sMMO (i.e., *Mycobacterium* sp. 002887815, *Mycobacterium* sp. 003053865, *Mycobacterium chubuense* A, and *Mycobacterium rhodesiae A*; **Fig S8**). We note MAG STB_95 from this study was most closely related to *Mycobacterium* sp.007714185, which did not encode genes for methanotrophy (**Fig S8**).

The detection of putative methanotrophs in the *Mycobacteriaceae* merits additional context. Over the last 70 years, a handful of studies have sporadically provided direct and indirect evidence of methanotrophy in strains identified as members of the genus *Mycobacterium* (20–22). This conflicting evidence may explain why members of *Mycobacteriaceae* have been precluded from recent surveys of methanotrophic taxa (23, 24). Up until very recently, the research supporting methanotrophy in the *Mycobacteriaceae* conflicted with numerous mechanistic studies focussed on the alkane and alkene degradation in strains such as *Mycobacterium chubuense* NBB4 and *Mycobacterium* sp. TY-6, where methane oxidation did not occur (25–28). To the best of our knowledge, the recent cultivation of *Candidatus* Mycobacterium methanotrophicum from an acidic cave biofilm is the first to provide direct physiological evidence of aerobic methane oxidation via the sMMO complex in a member of the *Mycobacteriaceae* (29).

We note the recent discovery of *Candidatus* Mycobacterium methanotrophicum here because the phylogenetic analyses carried out as part of that study aligns with our own phylogenomic analyses identifying putative methanotrophs in the *Mycobacteriaceae*. In the original study detailing methanotrophy in *Candidatus* Mycobacterium methanotrophicum, the authors showed that multiple members of the *Mycobacteriaceae* including *Candidatus* Mycobacterium methanotrophicum carried *mmoX* and *mmoB* homologues (29). We identified several of these strains in our phylogenomic analyses as members of the *Mycobacteriaceae* harbouring genes required for a complete or near-complete sMMO complex (e.g., *Mycobacterium rhodesia* A corresponding to *Mycolicibacterium rhodesia* NBB3, *Mycobacterium chubuense* A corresponding to *Mycolicibacterium chubuense* NBB4, *Mycobacterium moriakense* corresponding to *Mycolicibacterium moriakense*, *Mycobacterium elephantis* corresponding to *Mycolicibacterium elephantis*, and *Mycobacterium pulveris* corresponding to *Mycolicibacterium pulveris*) (see **Fig S8**) (29).

We highlight these parallel observations to underscore how phylogenomic analyses can be used to challenge old metabolic paradigms. With several *Mycobacteriaceae* strains being available through culture collection, there is an opportunity to follow up on the recent evidence of methanotrophy in *Candidatus* Mycobacterium methanotrophicum to assess whether other members of the *Mycobacteriaceae* in different habitats have been overlooked with respect to their contributions to the global methane cycle.

### Supplementary Figures

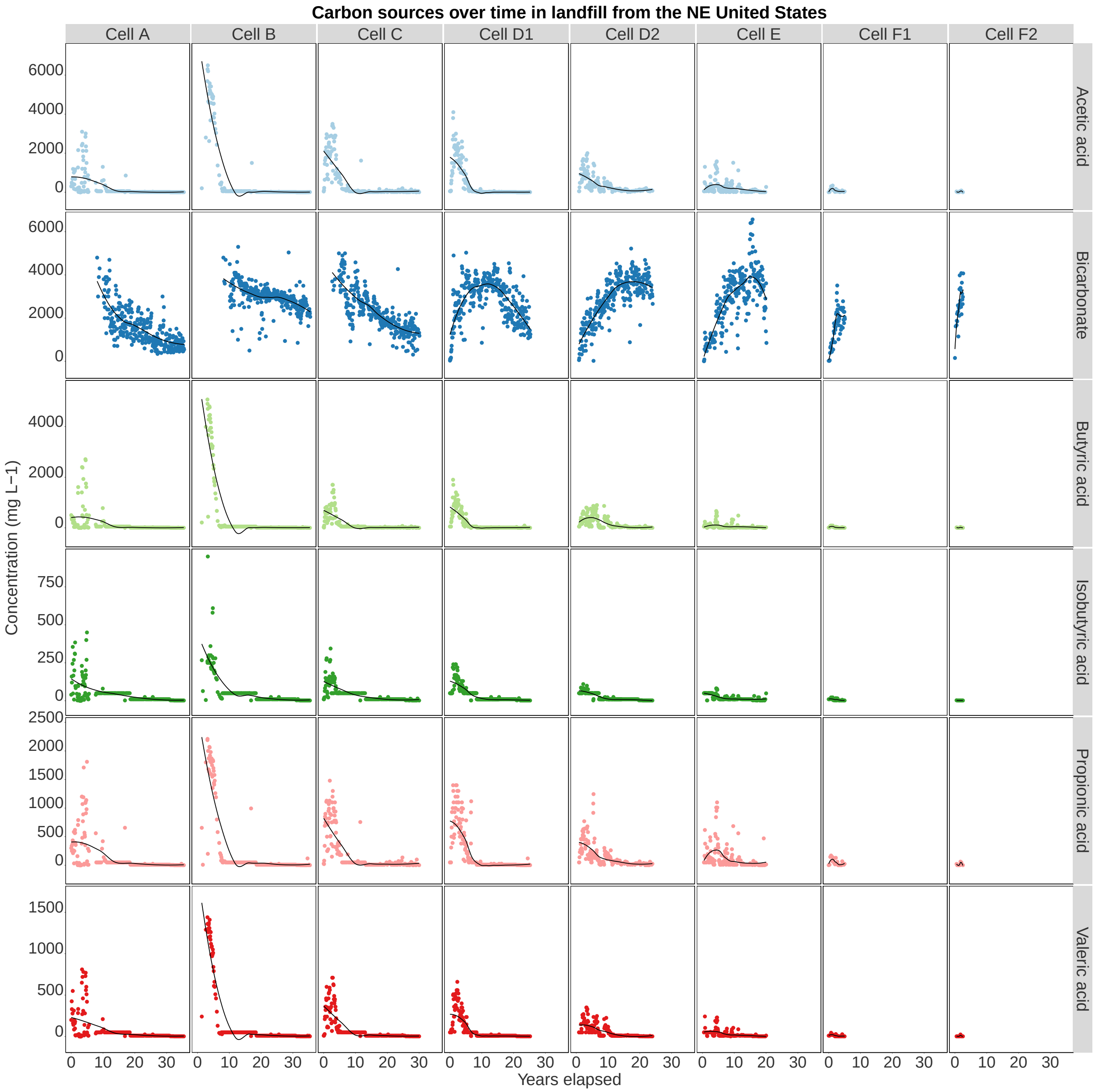


**Figure S1**: Organic acid and bicarbonate concentrations in landfill leachate collected from landfill cells A, B, C, D1, D2, E, F1 and F2 from 1983 to 2019. These data were compiled from monitoring records provided by the site management. Loess curves have been fitted to the data to visualize smoothed temporal trends.

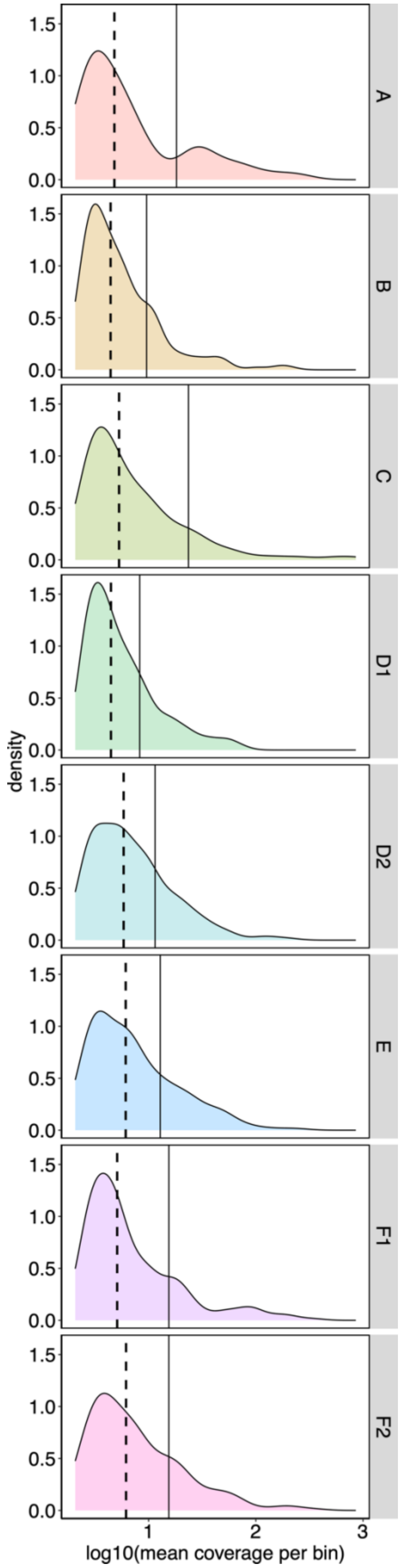


**Figure S2**: Density plots showing the distribution of mean coverage values calculated for all MAGs recovered from landfill cells A, B, C, D1, D2, E, F1, and F2. Solid black vertical lines denote the mean coverage calculated for MAGs at the whole-community level and dashed black lines denote the median coverage calculated for MAGs at the whole-community level.

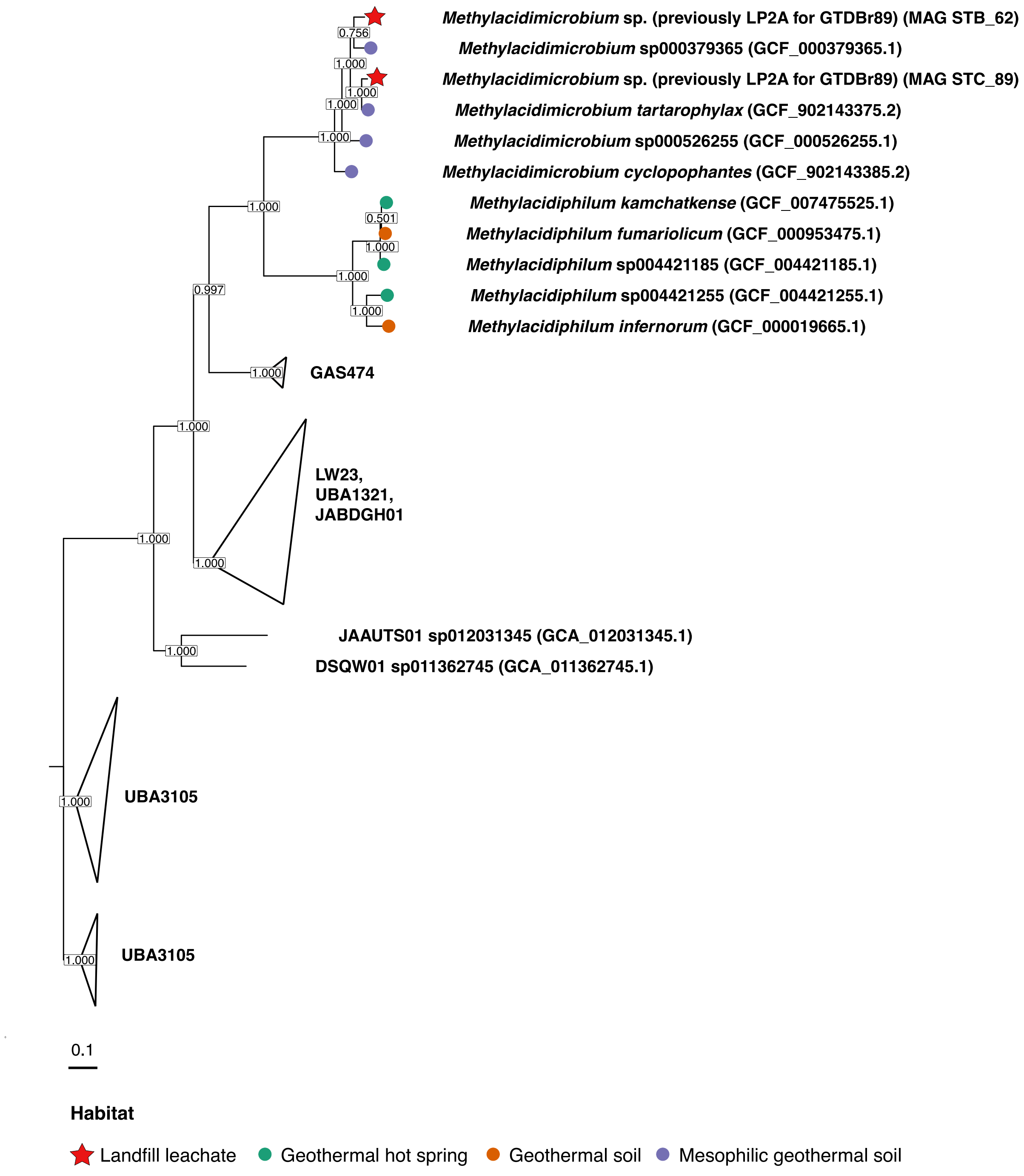


**Figure S3**: Rooted phylogenetic for *Methylacidiphilaceae* genomes retrieved from the landfill (denoted by a red star) and GTDB generated by GToTree. The unnamed family UBA3105 within the order *Methylacidiphilales* was used as the outgroup. Coloured circles correspond to the different environmental sources associated with the NCBI biosample entries for each genome. The scale bar denotes amino acid substitutions per site.

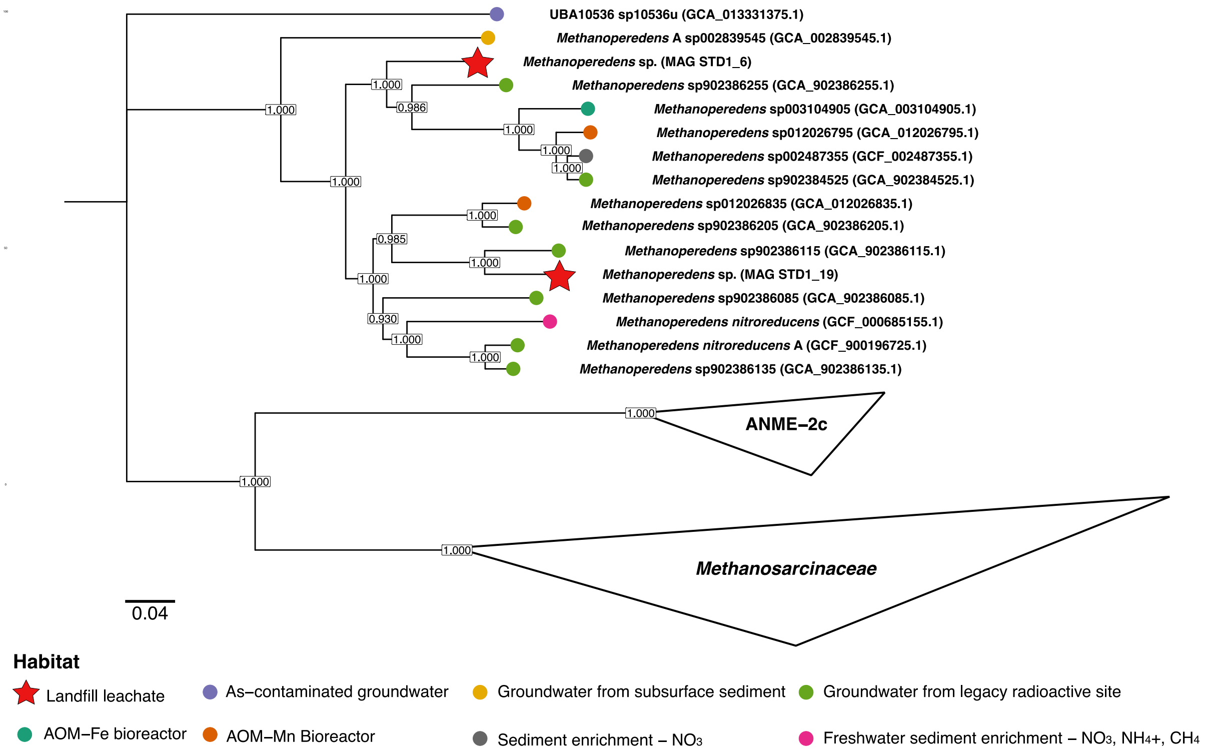


**Figure S4:** Rooted phylogenetic tree for *Methanoperedenaceae* genomes retrieved from the landfill (denoted by a red star) and GTDB generated by GToTree. The families ANME-2c and *Methanosarcinaceae* were used as outgroups. Coloured circles correspond to the different environmental sources associated with the NCBI biosample entries for each genome. The scale bar denotes amino acid substitutions per site.

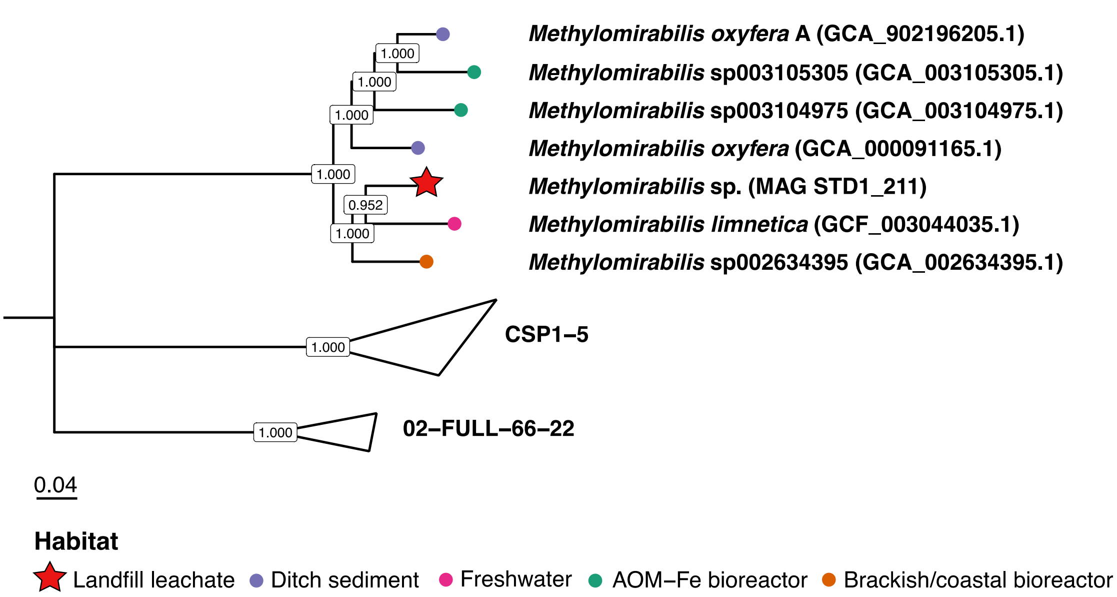


**Figure S5:** Rooted phylogenetic tree for *Methylomirabilaceae* genomes retrieved from the landfill (denoted by a red star) and GTDB generated by GToTree. The unnamed families CSP1-5 and 02-FULL-66-22 were used as outgroups. Coloured circles correspond to the different environmental sources associated with the NCBI biosample entries for each genome. The scale bar denotes amino acid substitutions per site.

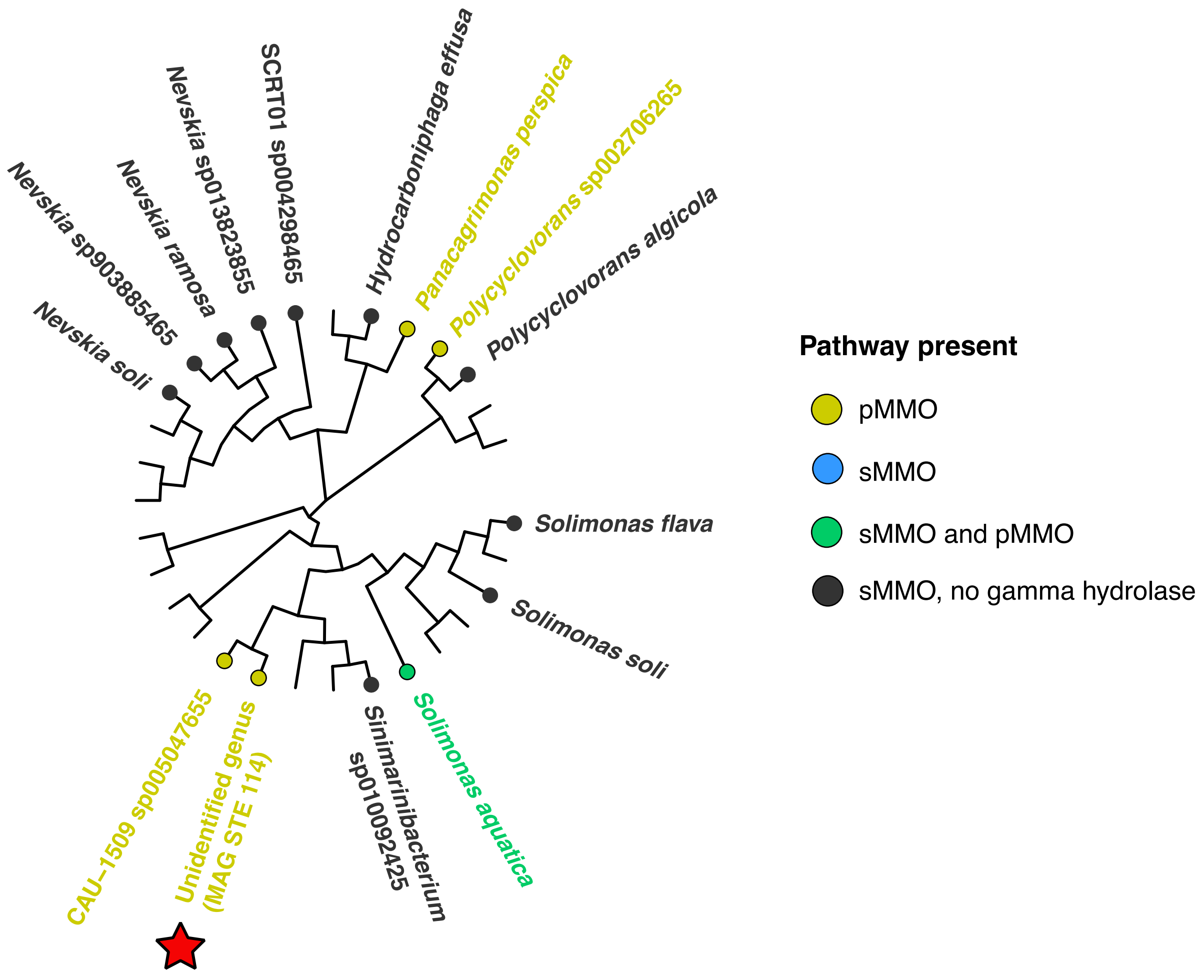


**Figure S6:** Unrooted phylogenetic tree for all 30 *Nevskiaceae* genomes retrieved from the landfill (shown with a red star) and GTDB, representing all currently available *Nevskiaceae* genomes. Coloured circles and text denote the different types of putative methanotrophs identified by including the Pfam entries for the pMMO and sMMO pathways during the tree building process in GtoTree.

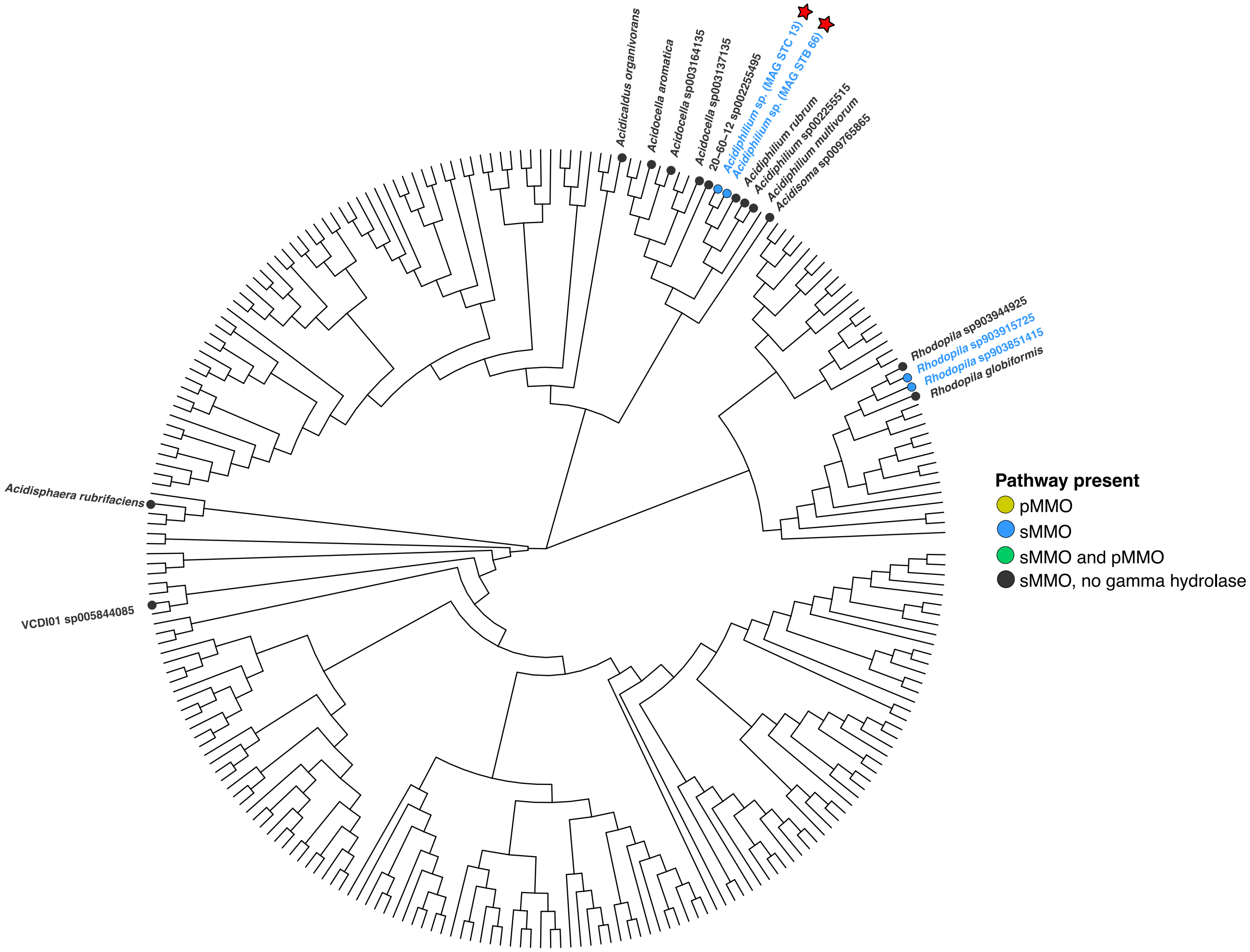


**Figure S7:** Unrooted phylogenetic tree for all 254 *Acetobacteraceae* genomes retrieved from the landfill (shown with a red star) and GTDB, representing all currently available *Acetobacteraceae* genomes. Coloured circles and text denote the different types of putative methanotrophs identified by including the Pfam entries for the pMMO and sMMO pathways during the tree building process in GtoTree.

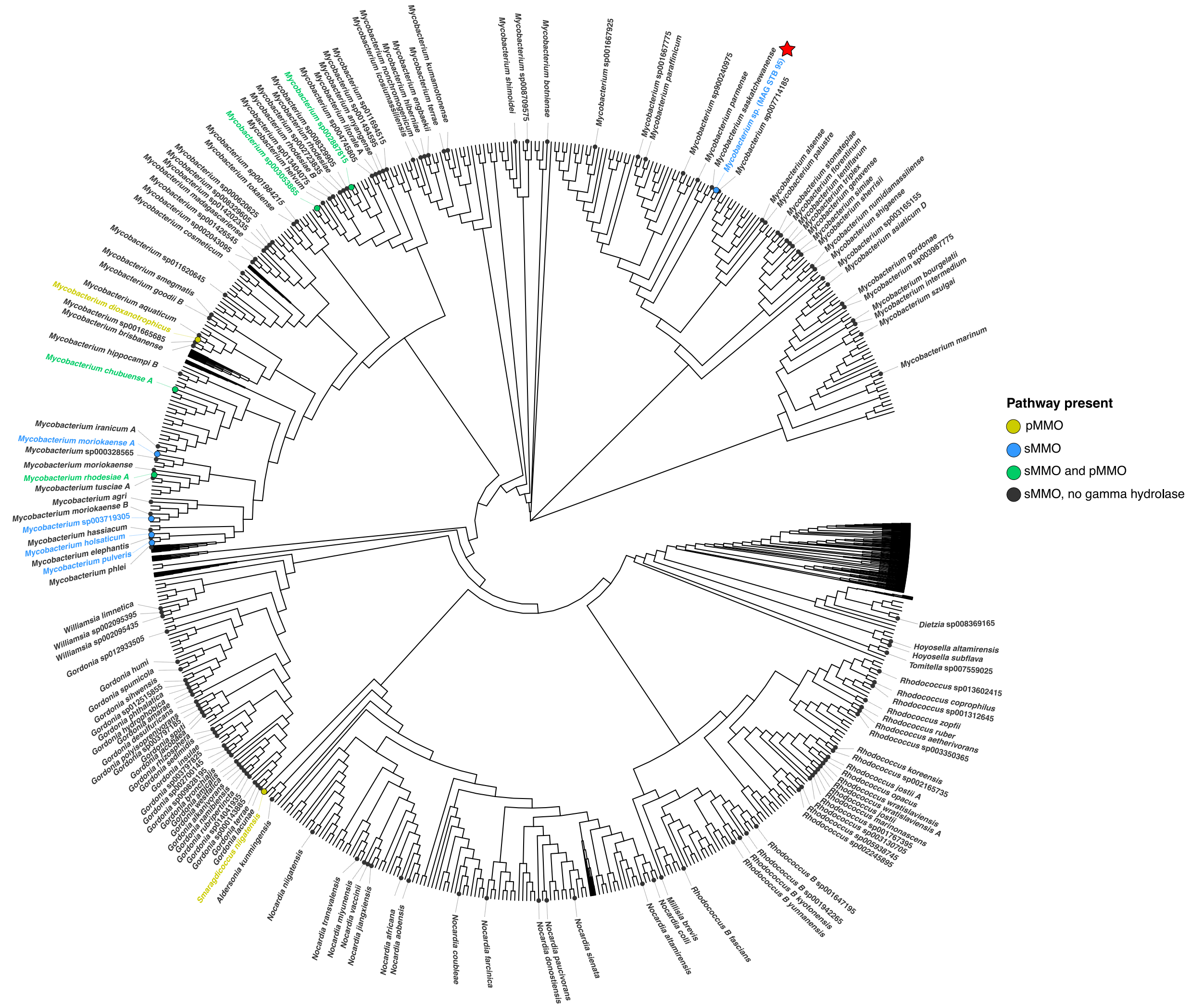


**Figure S8:** Unrooted phylogenetic tree for *Mycobacteriaceae* genomes retrieved from the landfill (shown with a red star) and GTDB. Coloured circles and text denote the different types of putative methanotrophs identified by including the Pfam entries for the pMMO and sMMO pathways during the tree building process in GtoTree. This tree includes all 811 representative genomes available through GTDB. Due to the size of the tree, clades with no hits for the categories of methane oxidation pathways present were scaled down to 10% of their size and hits for the pMMO, sMMO, and sMMO and pMMO categories had their symbols slightly increased in size.

**Table S1:** Summary statistics produced for all landfill metagenomes using BBMap ([https://sourceforge.net/projects/bbmap/](https://sourceforge.net/projects/bbmap/)).

| Site | # of scaffolds | # of contigs | Total scaffold length (bp) | Total contig length (bp) | Gap % | N50 for scaffolds | L50 for scaffolds | N50 for contigs | L50 for contigs | N90 for scaffolds | L90 for scaffolds | N90 for contigs | L90 for contigs | Maximum scaffold length (bp) | Maximum contig length (bp) | Sequence count for scaffolds >50 KB | Sequence size % for scaffolds >50 KB | Avg GC content | STDEV for GC content | Min. cov. for single MAG | Max. cov for single MAG | Mean cov for all MAGs | Median cov for all MAGs |
|---|---|---|---|---|---|---|---|---|---|---|---|---|---|---|---|---|---|---|---|---|---|---|---|
| A | 1.05E+06 | 1.06E+06 | 2.19E+09 | 2.19E+09 | 0.013 | 2.22E+05 | 2.06E+03 | 2.26E+05 | 2.04E+03 | 8.38E+05 | 1.10E+03 | 8.44E+05 | 1.10E+03 | 1.55E+06 | 1.55E+06 | 1.30E+03 | 6.65 | 0.50 | 0.11232 | 2.07 | 267.78 | 18.21 | 4.79 |
| B | 1.18E+06 | 1.19E+06 | 2.56E+09 | 2.56E+09 | 0.014 | 2.42E+05 | 2.20E+03 | 2.46E+05 | 2.19E+03 | 7.99E+05 | 1.21E+03 | 8.06E+05 | 1.20E+03 | 1.57E+06 | 1.57E+06 | 1.40E+03 | 6.36 | 0.47 | 0.10202 | 2.14 | 195.25 | 9.58 | 4.42 |
| C | 9.28E+05 | 9.41E+05 | 2.13E+09 | 2.13E+09 | 0.016 | 1.76E+05 | 2.43E+03 | 1.80E+05 | 2.40E+03 | 7.27E+05 | 1.13E+03 | 7.36E+05 | 1.13E+03 | 1.32E+06 | 1.10E+06 | 1.38E+03 | 7.29 | 0.48 | 0.12012 | 2.30 | 857.19 | 23.54 | 5.30 |
| D1 | 1.52E+06 | 1.53E+06 | 3.17E+09 | 3.16E+09 | 0.013 | 3.40E+05 | 2.08E+03 | 3.44E+05 | 2.07E+03 | 6.65E+05 | 1.49E+03 | 6.72E+05 | 1.48E+03 | 1.17E+06 | 1.17E+06 | 1.32E+03 | 4.58 | 0.46 | 0.10489 | 2.16 | 76.15 | 8.25 | 4.45 |
| D2 | 1.07E+06 | 1.08E+06 | 2.73E+09 | 2.73E+09 | 0.022 | 1.74E+05 | 2.94E+03 | 1.79E+05 | 2.90E+03 | 5.74E+05 | 1.44E+03 | 5.83E+05 | 1.44E+03 | 7.84E+05 | 5.33E+05 | 2.31E+03 | 7.95 | 0.43 | 0.10683 | 2.24 | 205.95 | 11.49 | 5.85 |
| E | 8.14E+05 | 8.21E+05 | 1.95E+09 | 1.95E+09 | 0.02 | 1.49E+05 | 2.61E+03 | 1.52E+05 | 2.59E+03 | 6.33E+05 | 1.15E+03 | 6.38E+05 | 1.15E+03 | 7.14E+05 | 6.63E+05 | 1.16E+03 | 5.56 | 0.46 | 0.11564 | 2.26 | 236.45 | 12.84 | 6.11 |
| F1 | 1.06E+06 | 1.07E+06 | 2.51E+09 | 2.51E+09 | 0.022 | 2.02E+05 | 2.60E+03 | 2.07E+05 | 2.57E+03 | 7.32E+05 | 1.23E+03 | 7.39E+05 | 1.23E+03 | 1.24E+06 | 1.10E+06 | 1.29E+03 | 4.70 | 0.45 | 0.10868 | 2.08 | 353.67 | 15.44 | 5.09 |
| F2 | 9.50E+05 | 9.61E+05 | 2.29E+09 | 2.29E+09 | 0.021 | 1.77E+05 | 2.66E+03 | 1.81E+05 | 2.63E+03 | 7.37E+05 | 1.16E+03 | 7.45E+05 | 1.15E+03 | 7.35E+05 | 7.35E+05 | 1.30E+03 | 4.93 | 0.43 | 0.09613 | 2.12 | 347.40 | 15.43 | 6.16 |
